# ‘Radical’ differences between two FLIM microscopes affect interpretation of cell signaling dynamics

**DOI:** 10.1101/2023.08.18.553714

**Authors:** Sravasti Mukherjee, Jeffrey Klarenbeek, Farid El Oualid, Bram van den Broek, Kees Jalink

**Affiliations:** Dept. of Cell Biology, The Netherlands Cancer Institute, Plesmanlaan 121, 1066CX Amsterdam; Swammerdam Institute of Life Sciences, University of Amsterdam, Science Park 904, 1098 XH Amsterdam; UbiQ Bio B.V., Science Park 301, 1098 XH, Amsterdam; BioImaging facility, The Netherlands Cancer Institute, Plesmanlaan 121, 1066CX Amsterdam

## Abstract

Emerging evidence suggests that cell signaling outcomes depend not only on the signal strength but also on its temporal progression. Our lab employs Fluorescence Lifetime Imaging of Resonance Energy Transfer (FLIM/FRET) biosensors to study intracellular signaling dynamics. We studied activation of β1 receptors by Isoproterenol, which triggers cAMP signaling via the G protein G_αs_, using two different FLIM microscopes: a widefield frequency domain FLIM (fdFLIM) setup and a fast confocal Time Correlated Single Photon Counting (TCSPC) setup.

When comparing results from each FLIM setup, unexpectedly we obtained distinctively different cAMP kinetics: fdFLIM recording of cAMP in HeLa and Cos7 cells yielded transient responses, reminiscent of rapid receptor desensitization, while TCSPC recordings exhibited sustained responses lasting over 30 minutes. We initially suspected phototoxicity due to the intense light locally in the laser focus spot in confocal microscopy to interfere with normal termination of signal transduction and set out to map photosensitive steps in the signaling cascade in detail. We find no evidence for light-sensitivity in either generation or breakdown of cAMP, but rather, our findings show that the kinetic differences are due to selective degradation of β1 agonists on the fdFLIM setup. Agonist degradation appeared due to the commercial FluoroBrite medium, even though this has been specifically advertised to lower phototoxicity and reduce autofluorescence. Mass spectrometry identified Folic acid, an undisclosed constituent of FluoroBrite, as the culprit leading to artifacts in fdFLIM measurements.

In all, our study underscores the impact of subtle phototoxicity effects on experimental outcome, and it shows that in this case confocal TCSPC provides the more reliable data needed to study response kinetics. This work also emphasizes the it is crucial that scientific vendors fully disclose chemical formulations.

## INTRODUCTION

The function and fate of the various cell types that make up tissues are tightly controlled by a large number of extracellular messenger molecules such as hormones, growth factors and cytokines. Cell surface receptors at the plasma membrane pick up the extracellular signals and transduce them into intracellular messages, so-called 2^nd^ messengers of which a few dozen have been identified to date. How can such a large number of different signals -well over a 1000 cell surface receptors have been identified-be relayed with just a limited set of 2^nd^ messenger signals? It has become increasingly clear that the answer, at least in part, lies in the tight spatial and temporal control of the ensuing intracellular signals^1–4^.

To fully appreciate the intricacies of spatiotemporal patterning of intracellular messages, these processes have to be studied with methods that provide both high spatial as well as temporal resolution, in individual living cells, and in a quantitative manner, i.e. by live cell fluorescence microscopy. Over the last decades, fluorescence microscopes have become faster, more sensitive, more photon-efficient and more versatile, whereas modern software allows real-time segmentation of cells and quantification of the results. To match the power of these microscopes, hundreds of fluorescent sensors were developed and optimized. These allow imaging all major second messengers: ions, such as Ca^2+^, Mg^2+^, Zn^2+^, Na^+^ and H^+^, small molecules like inositol trisphosphate (IP_3_), nitric oxide (NO), and cyclic adenosine monophosphate (cAMP), lipids like phosphatidyl inositol bisphosphate (PIP_2_), phosphatidyl inositol trisphosphate (PIP_3_) and diacyl glycerol (DAG), but also protein modifications such as phosphorylation and complex formation. The majority of these sensors employ Fluorescence Resonance Energy Transfer (FRET) between a donor-and acceptor fluorophore for read out.

The most straightforward manner to quantify FRET is by imaging the excited state lifetime of the FRET donor using Fluorescence Lifetime Imaging Microscopy (FLIM). FLIM is independent of sensor bleaching and concentration, light source fluctuations and a range of microscope imperfections that hamper quantification in intensity measurements^5^. Moreover, recent developments have rendered FLIM microscopes fast enough to follow cell signals with sub-second time resolution. The two most common implementations in FLIM microscopy are frequency-domain (fdFLIM) instruments and time-correlated single photon counting (TCSPC) instruments.

fdFLIM is usually implemented on widefield microscopes equipped with a fast gated camera. Our Lambert Instruments widefield fdFLIM system detects the delay between 40 MHz sine wave-modulated excitation light and the ensuing modulated fluorescence emitted from the cells. The latter is retrieved by recording a stack (a minimum of 3 but typically 12) of emission images at different relative phase delays with respect to the excitation^6–8^, using a phase-sensitive camera system. To minimize excitation, we introduced a new CCD camera with on-board phase demodulation^9^ which doubled photon efficiency. Typical excitation powers, at the plane of the cells, are as low as 400-900 µW (0.2-0.5W/cm^2^), well within the range of low-exposure timelapse experiments (e.g. Ca^2+^ imaging) seen in literature^10^.

In contrast, TCSPC is typically implemented on confocal point-scanning microscopes. Here, fluorophores in the confocal illumination spot are excited with sub-nanosecond laser pulses and the delay between laser pulse and arrival of ensuing emission photons at the detector is calculated by fast electronics. Until recently TCSPC was restricted to low photon count rates (< ∼10^6^/s) and collection of 515×512 pixel images required photon accumulation for up to several minutes; thus fast lifetime imaging was the domain of fdFLIM instruments. In a collaboration with Leica, we tested and further optimized a high-speed instrument (the Fast Lifetime Contrast system, FALCON) capable of acquiring good quality full-frame FLIM images every one or two seconds. After these optimizations, both setups have been routinely in use in our lab to record series of thousands of FLIM images without appreciable bleaching or obvious phototoxic effects.

While extremely powerful, fluorescence microscopy is not without its shortcomings. One major limitation is that intense excitation typically causes the dyes and sensors to bleach, thereby decreasing signal-to-noise over the time course of the experiment. In addition, excitation -in particular at the blue part of the spectrum-may be toxic to the cells and/or may affect various aspects of cell physiology^11–13^. In particular phototoxicity may vary strongly non-linearly with excitation intensity^13^, and because in confocal laser scanning microscopy excitation is restricted to a single focal laser spot, widefield microscopy is often considered to be milder for the cells^14^. In addition, the influence of cell culture media on light-induced cytotoxicity and bleaching has been recognized for decades^15–17^, leading to the formulation of e.g. Riboflavin-free media aimed at mitigating both cytotoxicity and bleaching.

Our lab studies dynamic aspects of agonist-induced cAMP changes using quantitative FLIM microscopy^18,19^ using an EPAC-based sensor that is specifically optimized for FLIM detection^20^. These studies require long timelapse imaging and therefore we had taken every effort to minimize excitation intensity. While comparing results obtained with our widefield fdFLIM setup (Lambert Instruments dedicated ultra-efficient FLIM camera)^6,9^ attached to a Leica DMI IRE2 to those obtained with fast TCSPC instruments (Leica FALCON on SP8/ Stellaris 8 systems^19^), we unexpectedly observed that both the amplitude and the time course of agonist-induced increases in cAMP differ radically between those two setups. As for our studies it is crucial that signal transduction dynamics are recorded faithfully, the present study was undertaken to illuminate the underlying cause of these differences.

We hypothesized and investigated several molecular mechanisms, including that intense laser excitation might interfere (directly or indirectly) with enzymes that control cAMP turnover. Contrary to expectation, we found that in this case TCSPC is the more reliable recording technique. Rather, we identify compounds present in the FluoroBrite imaging medium as the culprit. As most laboratories may not have routine access to both types of FLIM microscopes, our study also serves to caution microscopists not to blindly rely on the outcome of live cell imaging experiments, and to insist on knowing the molecular composition of all buffers and compounds used.

## RESULTS AND DISCUSSION

### Identical experiments yield widely different outcome depending on the microscopy setup

When carrying out time lapse experiments on two different FLIM microscopes in our lab, a Leica FALCON confocal TCSPC system and a Lambert Instruments widefield fdFLIM system, we routinely observed that these systems produced quite different results. In particular, the apparent rise in cAMP levels following stimulation of β1 receptors with Isoproterenol (IsoP) were transient (Fig. 1; Fig. S1A,C) on the fdFLIM setup whereas cells imaged on the confocal setup showed much more sustained cAMP responses which eventually declined towards baseline in > 60 min. As during these comparisons, we ascertained that on-stage cell culture conditions (imaging medium, temperature, CO_2_, and moisture conditions) were equal, differences in the microscopy setup were likely the culprit. Compared to widefield fluorescence microscopes, confocal point-scanning microscopes are often considered to be more detrimental as a result of the focused laser irradiation^14^ and it is thus tempting to speculate that the confocal setup might be introducing an artificially prolonged response as a result of phototoxicity.

**Figure 1:**
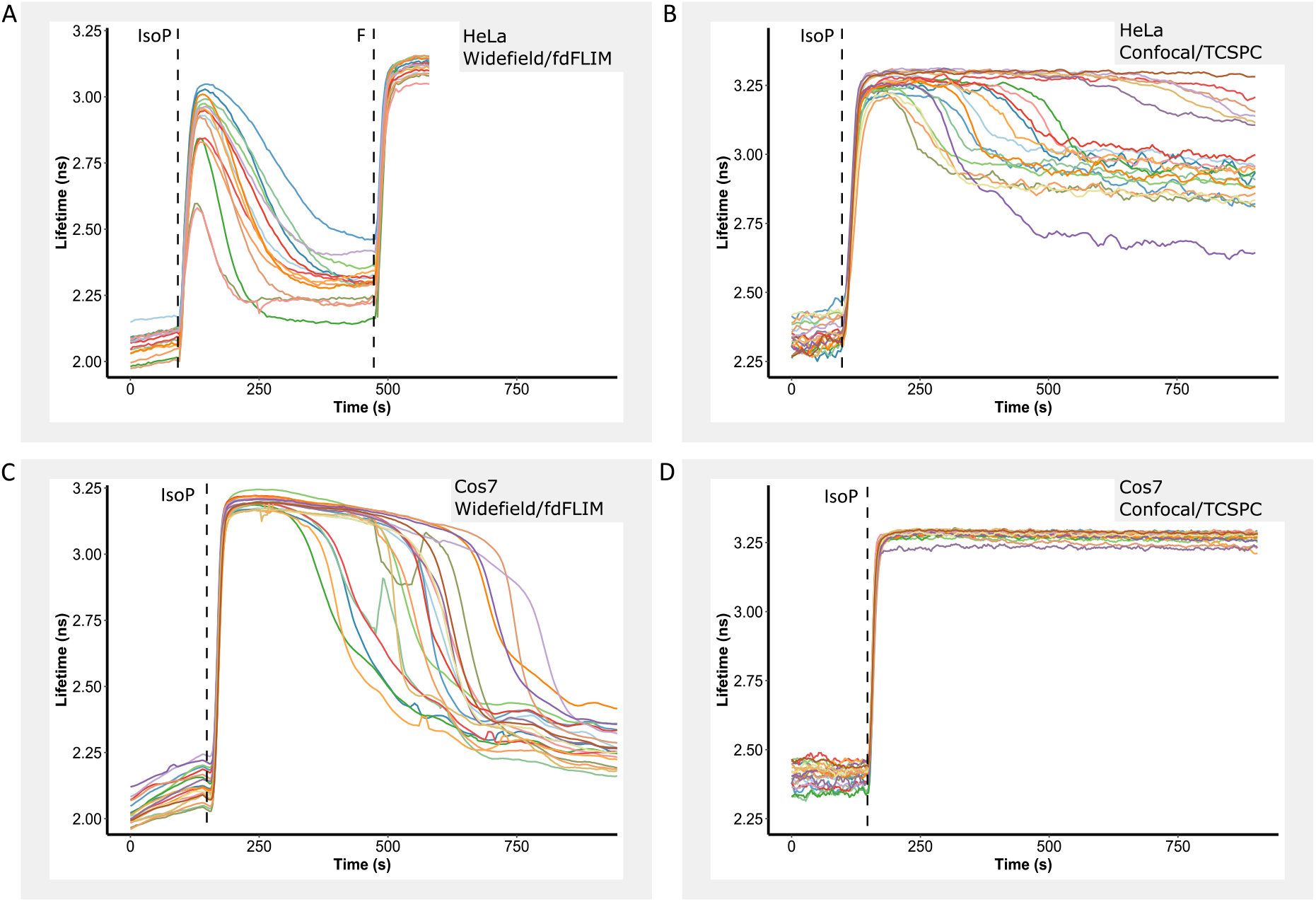
Identical experiments yield widely different response kinetics on two FLIM setups. Cells stably expressing the cAMP FLIM sensor Epac-S^H201^ or Epac-S^H250^ were imaged every 5 s. Following recording of a baseline, cells were challenged with Isoproterenol (IsoP; 40 nM) as indicated, and Forskolin (F; 25 µM) was added to saturate the response for comparison. Shown are FLIM timelapse responses in individual HeLa cells as detected on the **(A)** widefield fdFLIM setup and **(B)** on the FALCON TCSPC setup and in individual Cos7 cells detected on the **(C)** widefield fdFLIM setup and **(D)** on the FALCON TCSPC system. Note that the response to Isoproterenol as detected by fdFLIM did not saturate the FRET sensor and declined towards baseline within minutes (FWHM 110 +/- 50 s), whereas it did saturate the response on the FALCON setup, leading to sustained responses. Panels show a randomly selected subset of 20 cells taken from a single experiment. Full data sets (3 independent replicate experiment, 20 to 300 cells per experiment) are included in the supplemental material (Fig S1A,B,C,D)

To test this hypothesis, after recording a baseline and IsoP-induced transient response within a given field of view (FOV) we rapidly shifted to a few adjacent FOVs to check if the entire well was behaving uniformly (Fig. 2). Two independent experiments were performed on each set up. On the confocal setup, cells in all FOVs showed a similar sustained response. However, on the fdFLIM setup, cells in the original FOV had started to return towards their baseline cAMP levels whereas adjacent FOVs had maintained high cAMP levels and only upon imaging started to return to baseline after a few minutes.

**Figure 2:**
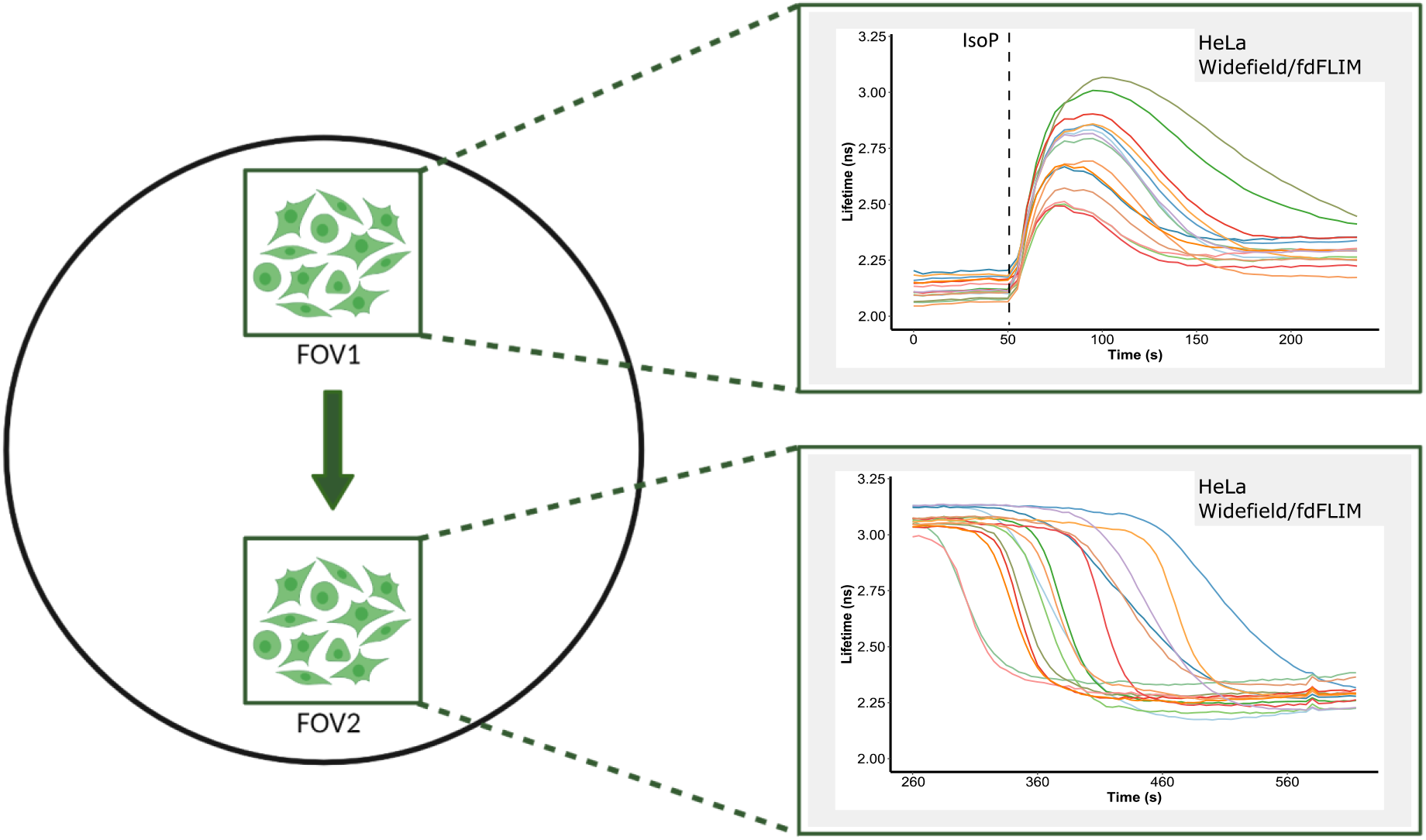
Field-shift experiment carried out on the widefield FLIM setup. Isoproterenol responses were recorded as in Fig. 1. Schematic representation of the experiment done in HeLa cells. After recording the response to Isoproterenol in a given field (FOV1), the stage was shifted to a second field (FOV2) and after rapid refocusing, recording was restarted. Similar effects were found in ∼300 cells in 3 independent experiments

These results indicated that one or more unknown factors on the widefield setup caused the rapid decrease in cAMP levels, but only in the FOVs that are exposed, suggesting that factors in the excitation regimen may play a role.

### Intensity and wavelength of the excitation light determine differences in cAMP dynamics

To explore this effect in more detail, we varied intensities of the excitation light on both setups. Note that we had previously optimized our fdFLIM measurements to allow long timelapse experiments by implementing a photon-efficient modulated charged coupled device (CCD) detector^6^, using on average only 0.9 mW of optical power at the FOV (0.5 W/cm^2^), at the low end of values typically used in widefield fluorescence microscopy. Nevertheless, further reduction of the intensity by 10-fold using a neutral density 1 (ND1) filter strikingly produced considerably longer cAMP responses (Fig. 3A-D).

**Figure 3:**
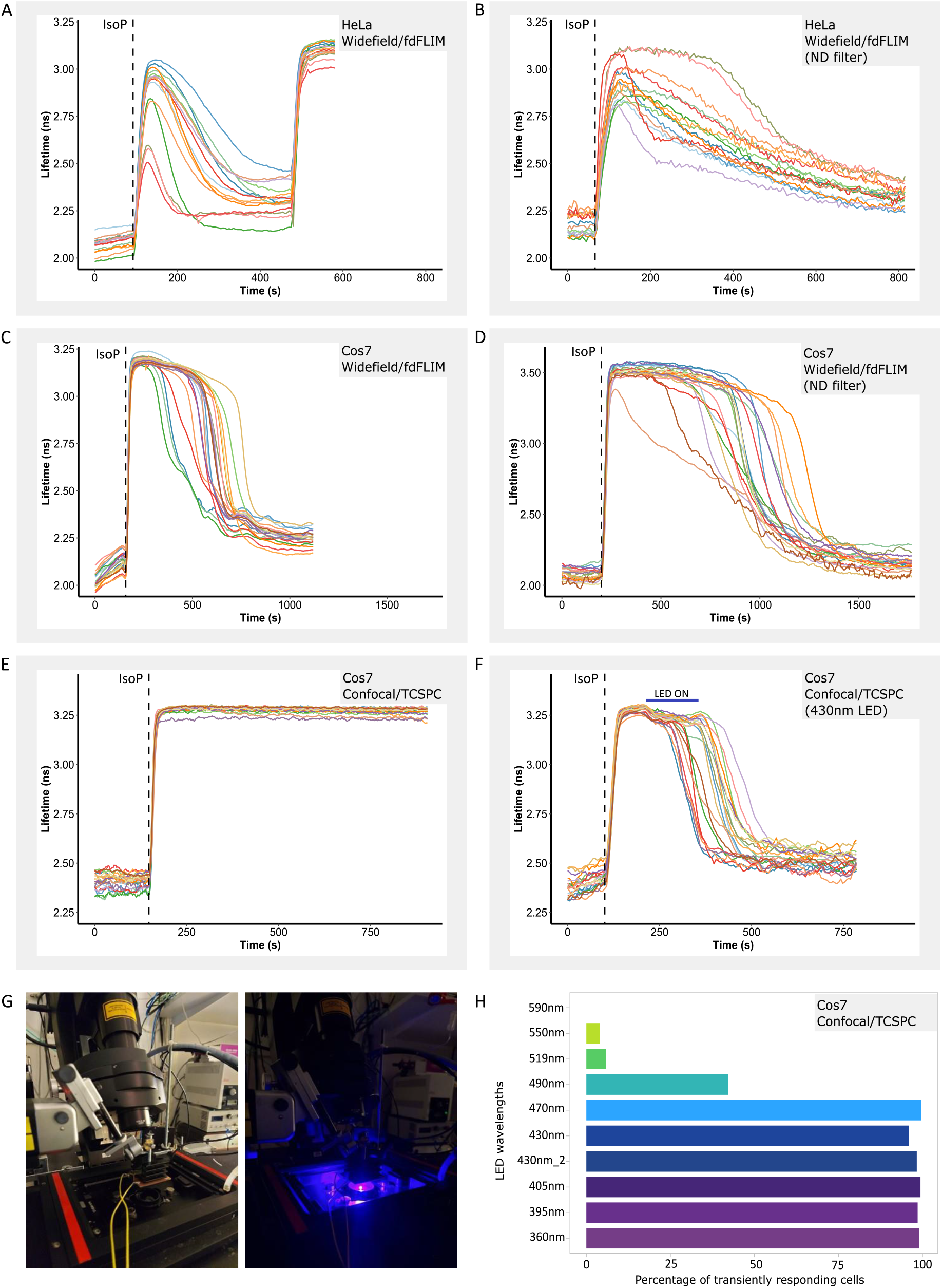
Blue light exposure affects the time course of cAMP dynamics. Responses of HeLa cells (A, B) and Cos7 cells (C-F) to 40 nM IsoP were recorded using fdFLIM (A-D) and using TCSPC (E,F). **(A)** Standard excitation at 0.9 mW (0.5 W/cm^2^) using a 435 nm sine-wave modulated LED. FWHM of the responses is 110 +/- 50 s. **(B)** Idem, using a ND1 filter in the excitation path to reduce excitation to 0.09 mW (0.05W/cm^2^) at the plane of the preparation. Response FWHM is 393 +/- 100 s. **(C, D)** Same experiments as in A and B for Cos7 cells; response FWHM were 337 +/- 50 s and 800 +/- 100 s, respectively. **(E)** Sustained responses detected by TCSPC at typical settings (45 µW; 7.5 mW/cm^2^). **(F)** Idem, except that cells were additionally exposed to continuous light from a 430 nm LED for 2 min as indicated. Note that this readily abrogated the sustained response (response FWHM 585 +/-200 s). **(G)** Image of the setup and LED. **(H)** Efficacy of LEDS with indicated wavelengths in abrogating sustained responses. Data depict % of cells with transient phenotype in 3 independent experiments. For A-F, representative traces were randomly selected from the data as detailed in M&M and Fig. 1.

Conversely, increasing the excitation power of the pulsed white light laser on the confocal microscope only marginally affected transientness, perhaps due to the limited power of the 448 nm laser line of the Leica white-light laser. We therefore increased exposure on the confocal system using a 430 nm power light emitting diode (LED) operated at 0.2 W/cm^2^ placed a distance of 8 mm above the cells (Fig. 3E-G). Indeed, exposure of the preparation to this light source for just 2 minutes sufficed to abrogate the sustained response (compare Fig. 3E to F). We also varied the color of the LEDs in these experiments (Fig. 3H) and found that wavelengths above 490 nm were ineffective.

Thus, we conclude that even modest excitation intensities of the order of 1 mW or less can induce an artificially rapid transient response in these experiments. In addition, our results indicate that in this case the confocal microscope, and not the wide field system, produces the more accurate results.

### Cell signaling, rather than the Epac FRET sensor, is affected by intense blue light

Theoretically, blue light might influence the dynamic FRET recordings in several manners. (i) It may affect the FRET reporter by changing either optical properties or cAMP binding to the Epac moiety; alternatively, blue light could induce true alterations in the time course of cAMP metabolism by (ii) augmenting the activity of PDEs, by (iii) inhibiting the action of ACs, or by (iv) attenuating upstream signaling components, e.g. by inducing receptor inactivation. These effects could either be direct, or secondary as a consequence of reactive oxygen species (ROS), which are notoriously formed under the influence of bright blue light^21,22^. We note that it is unlikely that blue light strongly affects the Epac sensor because after prolonged timelapse experiments, a rise in cAMP induced with Forskolin typically saturates the sensor (see Fig. 3A). Nevertheless we investigated this in an independent manner, namely by detecting cAMP levels biochemically by performing a cAMP ELISA on Cos7 cells expressing the Epac sensor as well as on WT cells not expressing the sensor.

The results in Fig. 4 show that in both cell lines, stimulation with 40 nM Isoproterenol readily increased cAMP levels, which was almost completely abolished upon exposure to 430 nm LED light for 2 minutes.

**Figure 4:**
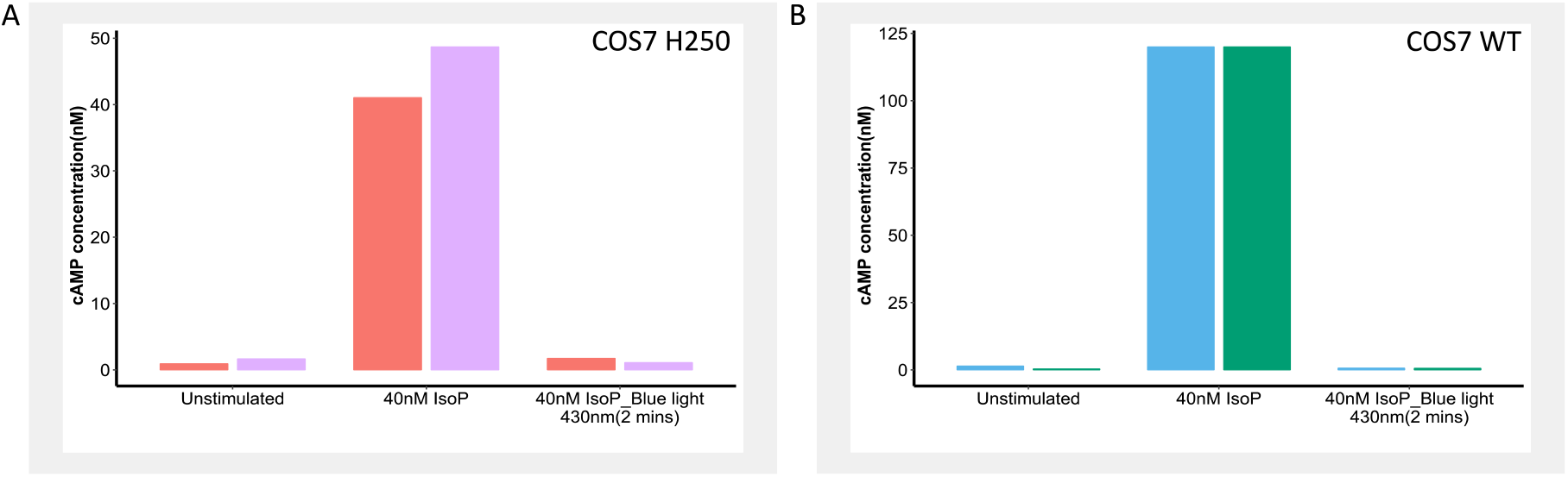
ELISA assays confirm that blue light affects cAMP levels. **(A)** cAMP levels under different conditions in Cos7 cells expressing the Epac sensor. Stimulation with 40 nM Isoproterenol raises cAMP levels, which is completely abolished by blue light (430 nm LED). **(B)** Same experiment as (A) repeated in WT Cos7 cells without the Epac sensor. Note that the response to 40 nM IsoP is an under-estimate as it maxed out the ELISA assay; see Materials and Methods. Bars represent biological duplicates.

This indicates that the transient response is not a sensor artifact but rather, that it truly affects response kinetics. We therefore next set out to discriminate between the other possibilities.

### High intensity blue excitation light generates Reactive Oxygen Species (ROS)

First, we sought to investigate whether blue-light induced ROS formation is involved, since ROS formation may affect biochemical reactions at various levels in the signal transduction cascade. Using cells loaded with the ROS dye (Di(Acetoxymethyl Ester) (6-Carboxy-2’,7’-Dichloro-dihydrofluorescein Diacetate)^23,24^, several FOVs were imaged at intensity settings commonly used on each of the setups, for 2 minutes. When scanning an overview region encompassing all imaged fields with the confocal, a dose-dependent dye staining was observed in all 3 FOVs imaged on the widefield system whereas no ROS production was observed in FOVs imaged with the confocal system (Fig. 5A). Thus, whereas confocal point scanning instruments are often considered to be more phototoxic, the opposite appears to be true when comparing wide field FLIM instruments to confocal TCSPC equipment. This is in line with our assessment that the confocal produces the more accurate temporal results in this case. As expected, exposure of cells to 430 nm LED illumination (compare Fig. 3F,G,H) on the confocal for 2 minutes also produced significant staining of the ROS dye (Fig. 5B, C). As expected, addition of the ROS scavenger Ascorbic acid (100 µM) to the imaging medium restored sustained responses completely (Fig. 5D), thus confirming that ROS formation is involved in inducing the transient cAMP phenotype.

**Figure 5:**
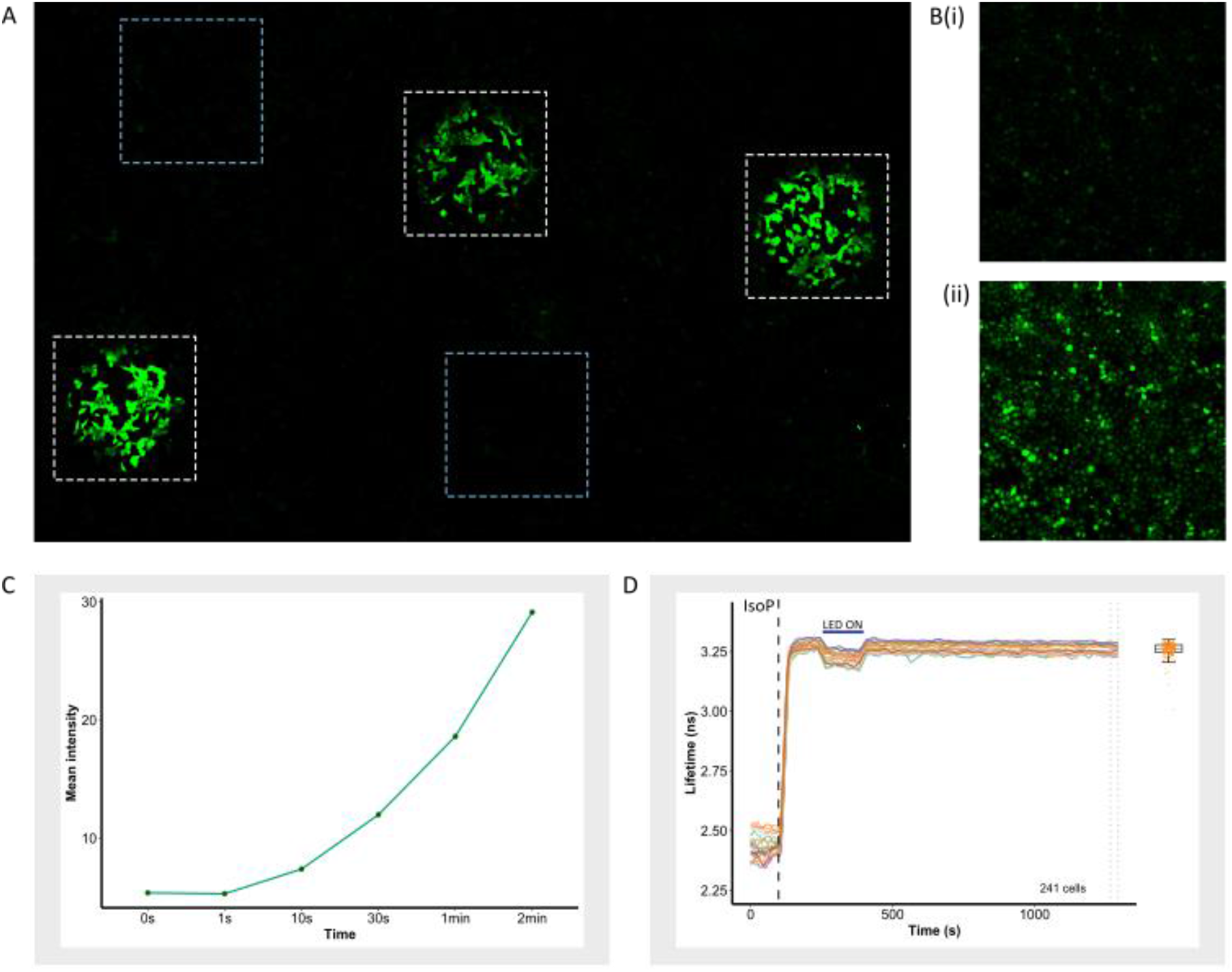
Widefield FLIM imaging evokes ROS formation. Confluent Cos7 cells were incubated with the ROS dye Di(Acetoxymethyl Ester) (6-Carboxy-2’,7’-Dichlorodihydrofluorescein Diacetate) and mounted on the microscope in FluoroBrite imaging medium. **(A)** Several fields were illuminated for 2 min on the fdFLIM setup (white dashed lines) and others on the TCSPC system (blue dashed lines). Note the absence of detectable ROS formation in FOV imaged on the confocal. **(B)** Formation of ROS induced by the 430 nm LED in the configuration of Fig. 3G **(i)** before shining 430 nm LED and **(ii)** 2min after shining 430 nm LED. **(C)** Time-dependence of ROS formation detected from dye intensity upon exposure to 430 nm LED for the indicated times. **(D)** Ascorbic acid reverts the cAMP response to sustained despite exposure to 430 nm LED light. B, C, D: Representative experiments are shown out of 2 repeats.

### Blue light does not alter cAMP turnover

Next we aimed to understand by what mechanism ROS might affect the kinetics of agonist-induced cAMP changes, in particular whether cAMP production by adenylate cyclases (ACs), or degradation through phosphodiesterases (PDEs) could be affected through ROS-mediated modification of the activity of those enzymes (hypotheses ii and iii; see above). In experiments aimed at illuminating the potential role of PDEs, we used UV-mediated photolysis (“uncaging”) of DMNB-caged cAMP, which induces fast increases in intracellular cAMP levels independent of membrane receptors and AC. In these experiments we observed no differences in the decay rate of cAMP levels with or without 470 nm LED illumination (Fig 6A).

**Figure 6:**
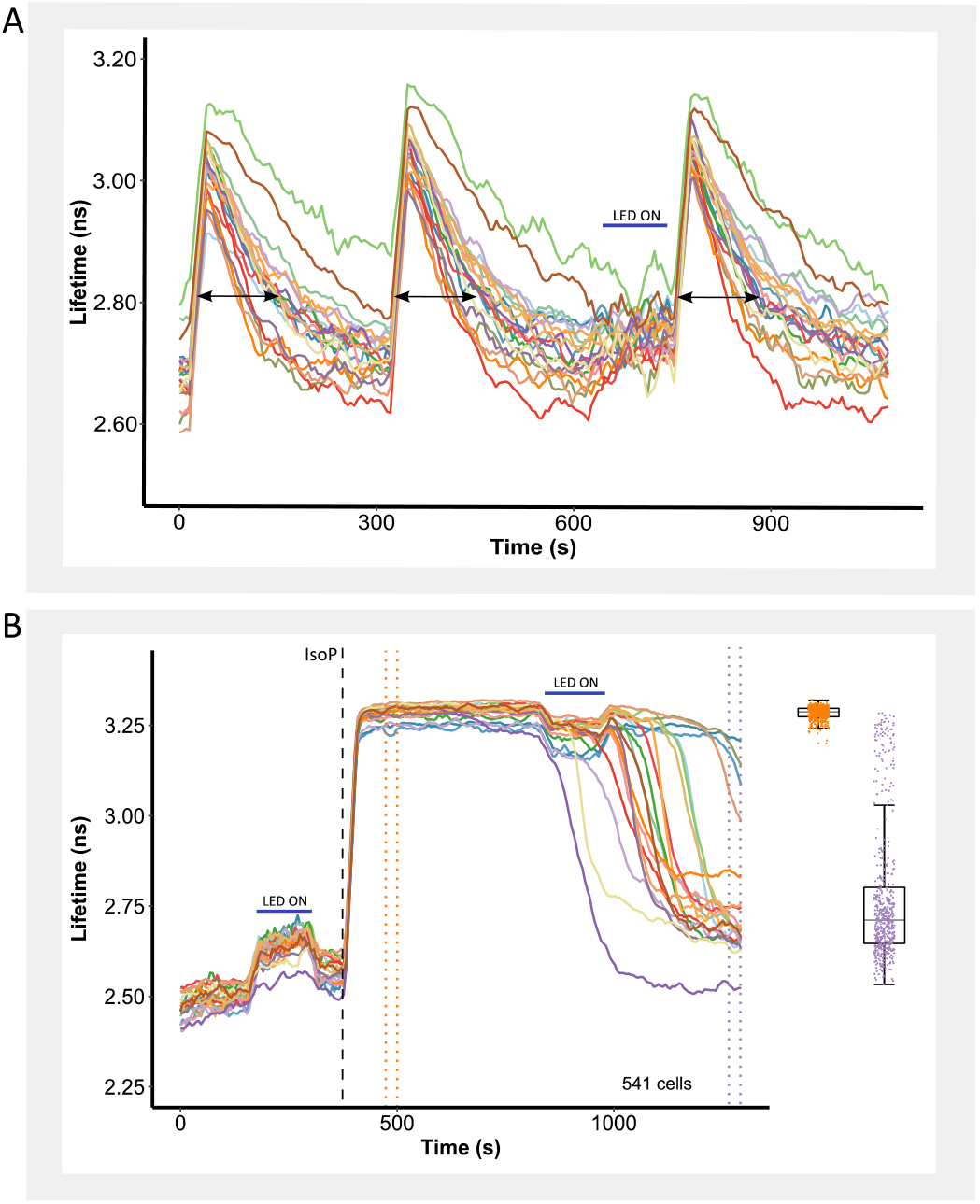
Blue light does not detectably alters generation or breakdown of cAMP. **(A)** Responses to three consecutive doses of cAMP generated by flash photolysis of caged cAMP. After the second dose, cells were exposed to LED light. cAMP decay times were identical (FWHM 23 +/- 5 s) for all 3 cases, indicating that PDE activity is unaltered by the light. Note that in this case 470nm LED was used to prevent slight photolysis of caged cAMP by the 430 nm LED. **(B)** Blue light fails to evoke response transientness when applied before addition of agonist. Shown are 20 randomly selected traces (see M&M) as well as the response magnitude of all 541 cells taken before (orange boxplot) and after (purple boxplot) LED exposure.

It is harder to directly address the opposite possibility, i.e. the downregulation by blue light/ROS of the activity of AC following G-protein coupled receptor (GPCR) stimulation. However, as it is apparent from the traces that the effects of blue light linger long after termination of the illumination (compare Fig. 3F), blue light should also attenuate the response when administered shortly before stimulation of the receptors. Surprisingly, however, we found that blue light only affected cAMP kinetics when administered *during* the cAMP response, and not *before* (Fig. 6B). Combined, these experiments indicate that neither modification of PDEs nor of ACs play a role in altering cAMP kinetics upon exposure to blue light/ROS. We thus focused our attention on more upstream events, i.e. at the level of the β-AR receptor or ligand binding to it.

### Blue light/ROS interferes in cAMP signaling up-stream

The surprising absence of effect when blue light was administered before initiation of GPCR signaling prompted us to focus on receptor-ligand interaction (hypothesis iv). First, we tested whether cAMP generated through other agonists and receptors was also affected by blue light. Indeed, the temporal responses to Norepinephrine (NE) and Adrenaline (AR), which both also activate β-adrenergic receptors, were similarly shortened in the presence of blue light (Fig. 7A-D). In contrast, however, the response to Prostaglandin-E1(PG), which acts through Prostanoid receptors was unaffected by exposure to blue light (Fig. 7E,F) Thus, the attenuating effect of blue light on the response might be restricted to β-receptor agonist or their interaction with the receptor.

**Figure 7:**
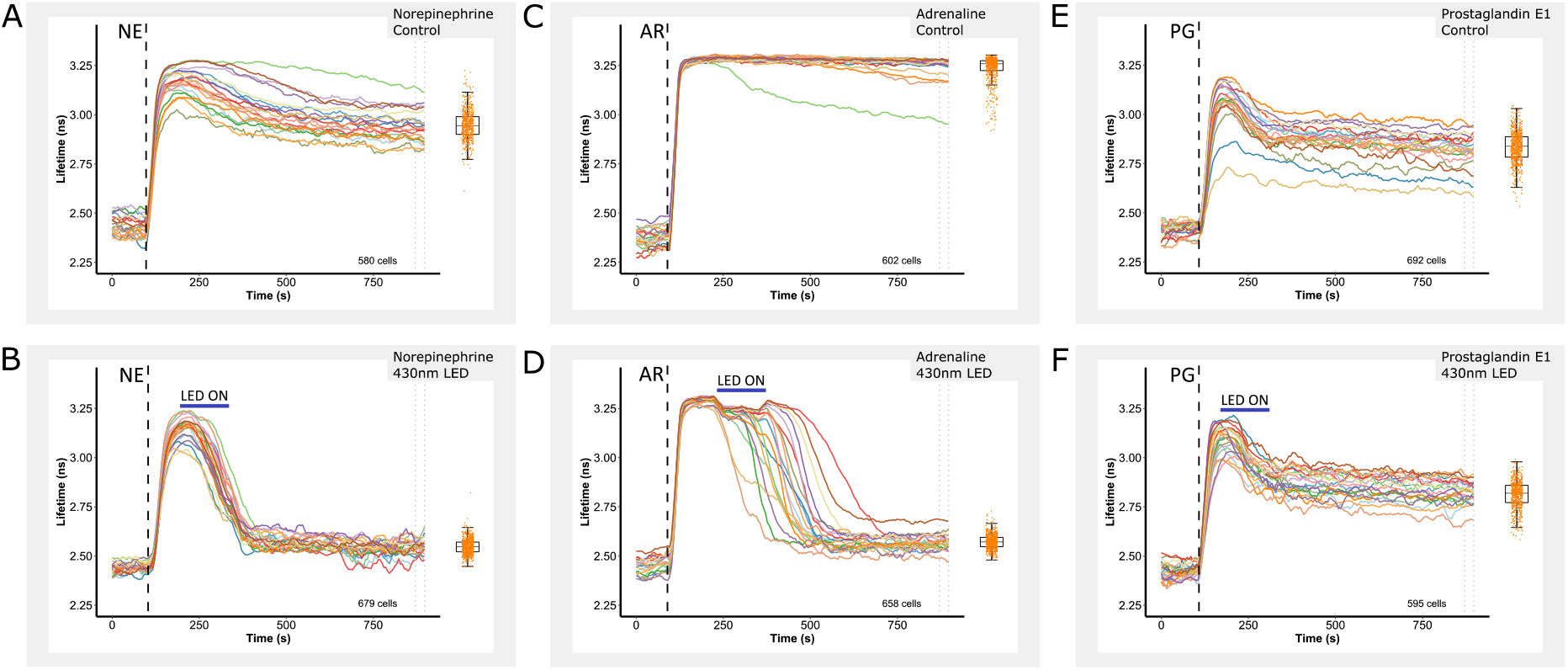
Blue light effects are agonist-and medium-specific. **(A)** Sustained responses of Cos7 cells to 200 nM Norepinephrine (NE) **(B)** Idem, altered to transient upon illumination with 430 nm LED light for 2 min. **(C)** Sustained responses to 250 nM Adrenaline (AR). **(D)** Idem, altered to transient upon exposure to 430 nm LED light for 2 min. **(E)** Typical, partially sustained responses of Cos7 cells to 200 nM Prostaglandin-E1 (PG). **(F)** Unaltered responses when Cos7 cells are stimulated with 200 nM Prostaglandin-E1 (PG) and illuminated with 430 nm LED light for 2 min. 20 randomly selected traces are displayed along with the quantification of all cells (boxplots) as detailed in M&M.

A literature search revealed three publications reporting that IsoP and NE might be degraded by strong blue light in cell culture media that contain elevated levels of Riboflavin (Vitamin B2), such as DMEM^25–27^. However, our experiments were carried out using FluoroBrite (FB)^28^, a specially formulated medium without Riboflavin^29^ which aims to reduce medium autofluorescence and protect cells against phototoxic effects during live-cell fluorescence microscopy. Nevertheless, we next set out to investigate whether interactions of the imaging medium could be responsible for the blue light-induced differential cAMP kinetics.

### FluoroBrite DMEM imaging medium causes photo-degradation of β-adrenergic agonists

To directly assess whether compounds in the FB imaging medium promote photo-degradation of the agonists, we exposed agonists dissolved in 200 µl of medium in eppendorf vials to 430nm blue light at an intensity comparable to our imaging experiments (Fig. 5-7) for 2 or 6 minutes. Strikingly, illumination strongly attenuated the potency of these agonists to induce cAMP responses in our reporter cells (Fig. 8A-C). This effect was fully rescued when ROS scavengers such as Ascorbic acid (100 µM) were included in the mix prior to exposure (Fig. 8D). Similar results were observed with Norepinephrine (NE; Fig. 8E,F) and Adrenaline (AR; Fig. 8G,H). In contrast, when Prostaglandin-E1 was added to FB medium and exposed to blue light, no attenuation of signaling was observed (PG; Fig. 9I,J).

**Figure 8:**
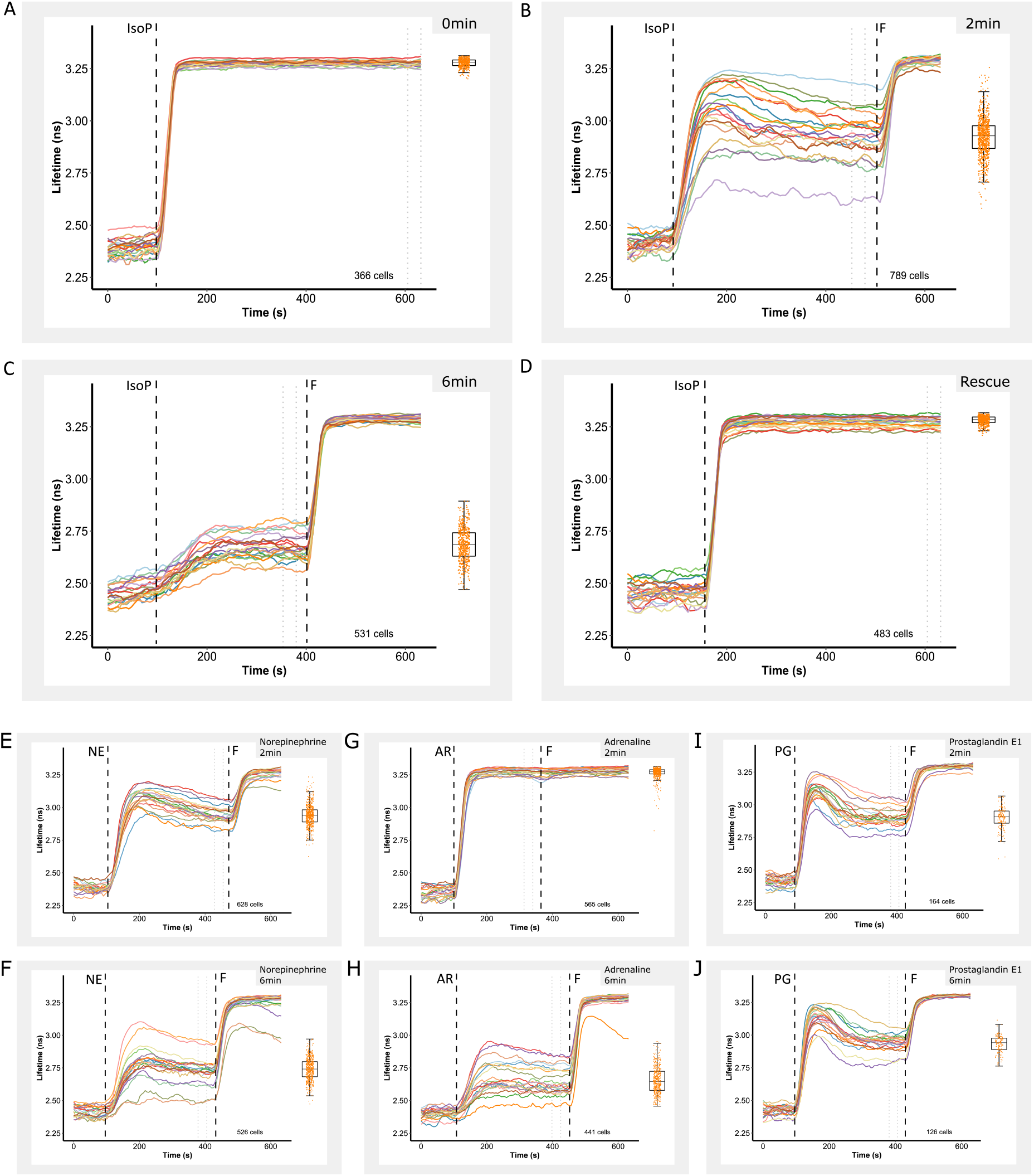
FluoroBrite medium mediates degradation of β-adrenergic agonists upon blue light illumination. **(A-D)** Responses to 40 nM Isoproterenol when added from small batches dissolved in 200 µl of FB medium that have been subjected to blue light illumination for (**A)** 0 min, **(B)** 2 min, **(C)** 6 min and **(D)** for 6 min in the presence of 100 µM Ascorbic acid. **(E, F)** Idem, responses to Norepinephrine (200 nM); shown are **(E)** after 2 min pre-exposure to blue light and **(F)** after 6 min pre-exposure to blue light. **(G,H)** Idem, responses to Adrenaline (250 nM); shown are **(G)** after 2 min pre-exposure to blue light and **(H)** after 6 min pre-exposure to blue light. **(I,J)** Idem, responses to Prostaglandin-E1 (200 nM) **(I)** after 2 min pre-exposure to blue light, and **(J)** after 6 min pre-exposure to blue light.

**Figure 9:**
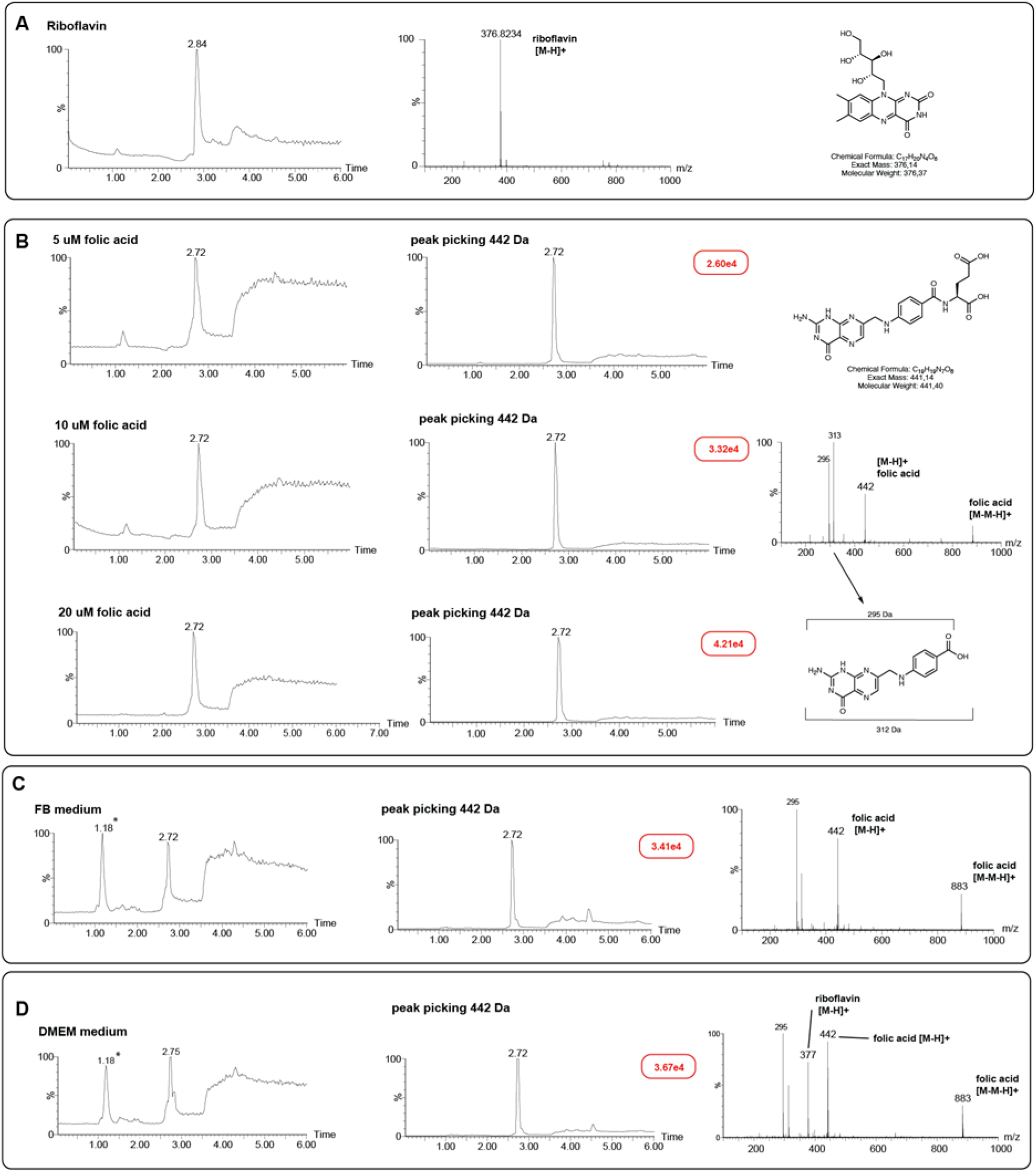
Mass spectrometry analysis of FluoroBrite medium shows absence of Riboflavin, but presence of Folic acid. **(A)** LC-MS analysis sample of pure Riboflavin in milliQ. (B) LC-MS analyses pure folic acid samples in milliQ at three concentrations. Note fragments with mass 295 Da and 313 Da (pteroic acid), formed by loss of the glutamic acid residue during analysis. (C) LC-MS analysis of FluoroBrite medium indicates presence of Folic acid but not Riboflavin. (D) LC-MS analysis of DMEM medium indicates presence of both Riboflavin and Folic acid. Based on the ion count (red boxes) of the three folic acid samples (see B) and that of the DMEM sample (see D), the concentration of folic acid in both DMEM and FB is around ±10 – 20 μM. The 1.18 min peak denoted with an asterisk in the chromatogram of FB (C) and DMEM medium (D) corresponds to polymeric material, as judged by the MS signal.

### Mass-Spectrometry analysis of FluoroBrite DMEM reveals presence of Folic Acid

Because the composition of FB medium is not disclosed on the webpages of Gibco, ThermoFisher^28^, we contacted the supplier to request this information. Unfortunately, access to those data was subject to signing of an extensive non-disclosure agreement. This is unacceptable because it would preclude publication of our findings, and we therefore had to embark on identifying possible photo-oxidative components in FB ourselves.

Samples of FB medium, hepes-buffered saline (HBS) and normal DMEM were compared by mass spectroscopy analysis and necessary controls of pure Riboflavin and Folic acid were taken along (Fig. 9A,B; See M&M). The results confirm absence of Riboflavin in FB medium (Fig. 9C). However, a conspicuous peak at 441 Da was common in both FB medium and normal DMEM which we tentatively identified as Folic acid by comparing the MW to the (fully disclosed) list of DMEM constituents^30^, the concentration of Folic acid in both mediums being in the range of 10-20 µM (Fig. 9C,D). Folic acid is a member of the Vitamin B family too (Vitamin B9) and it has been reported to degrade upon exposure to blue light^31,32^. However, no reports exist (to our knowledge) that in this process inactivation of bystander molecules such as Isoproterenol could take place.

To directly test this possibility, we repeated our pre-incubation experiments in HBS, except that FB medium was replaced with Folic acid at a concentration lying in the range of 10-20 µM (18 µM). Again, exposure to blue light for 2 or 6 minutes rendered the mix inactive in eliciting cAMP signals in reporter cells (Fig. 10A-C) and inclusion of 100 µM Ascorbic acid could completely rescue potency of the agonist in bio-assays (Fig. 10D). The effect of Folic Acid was also checked on responses induced by NE, AR and PG (Fig. S3). In line with this observation, while exposure of reporter cells to blue light after stimulation with IsoP resulted in sustained cAMP signaling in HEPES-buffered salt solutions (HBS) devoid of Folic acid (Figs. 10E), it caused a transient response in HBS spiked with 45 µM Folic acid (Fig. 10F). Again, inclusion of Ascorbic acid protected against IsoP inactivation (Fig. 10G). We conclude that 1) the presence of Folic acid in media such as FB causes unexpected artifacts when studying β-AR signaling, at least at conditions that involve exposure to blue light, and 2) that the refusal to disclose the composition of FB medium necessitated very time-consuming control experiments, thereby delaying our progress with several months.

**Figure 10:**
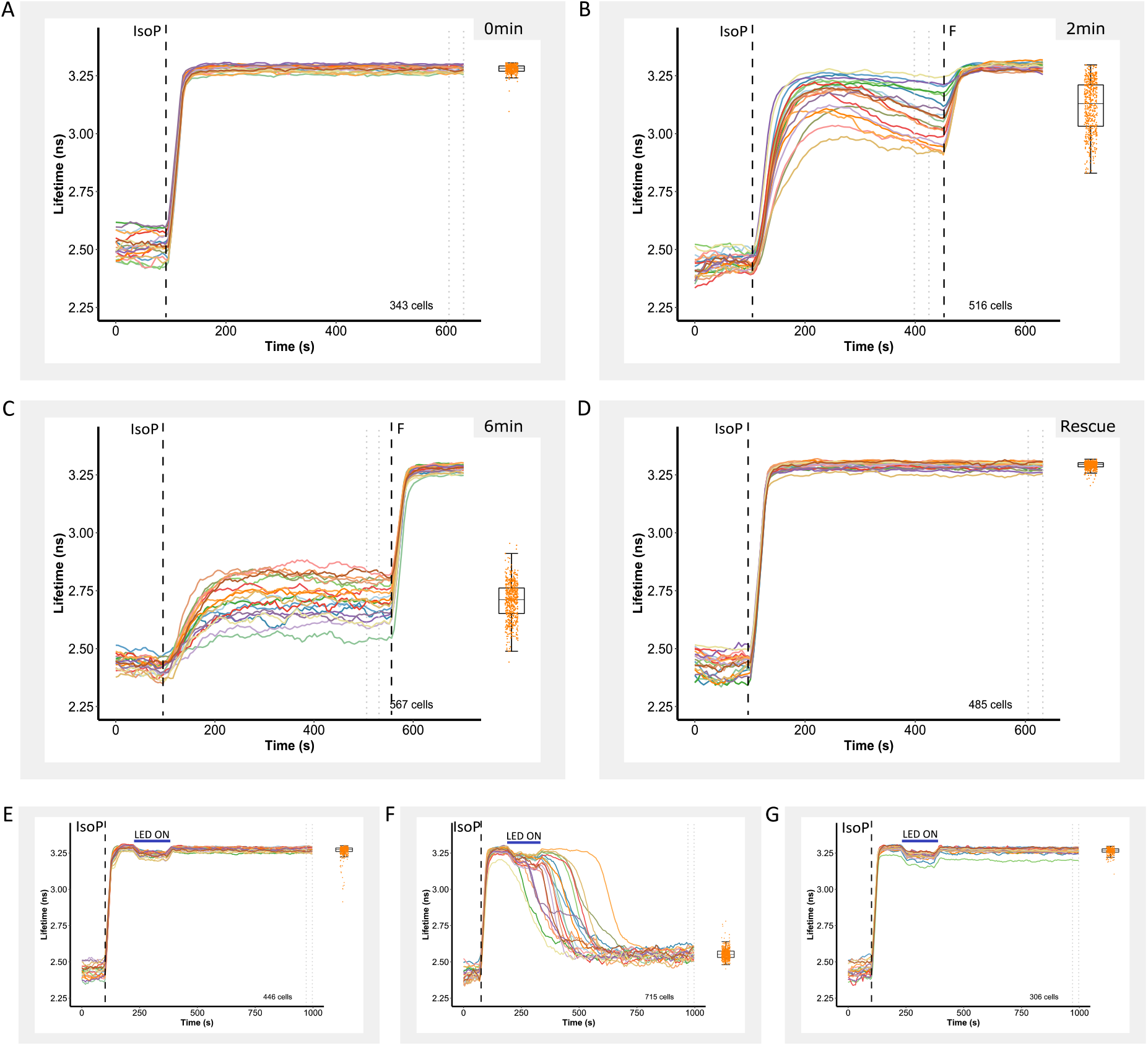
Folic acid present in FluoroBrite medium causes light-induced degradation of AR agonists. **(A,B,C)** Agonist (40 nM Isoproterenol) dissolved in 200 µl of HBS containing 18 µM Folic acid in eppendorf vials were subjected to blue light (430 nm LED) illumination for 0, 2, 6 min respectively before use in cell assays. **(D)** Agonist (Isoproterenol) dissolved in 200 µl of HBS containing 18 µM Folic acid in eppendorf vials along with 100 µM Ascorbic acid were subjected to blue light (430 nm LED) illumination for 6 min. Note protection of the agonist by Ascorbic acid. **(E)** Sustained responses of Cos7 cells in HBS medium only when stimulated with 40 nM Isoproterenol and illuminated with 430 nm LED for 2 min. **(F)** Transient responses of Cos7 cells in HBS medium containing 45 µM Folic acid when stimulated with 40 nM Isoproterenol and illuminated with 430 nm LED for 2 min. **(G)** Rescue of transient responses of Cos7 cells in HBS medium containing 45 µM Folic acid and 100 µM Ascorbic acid when stimulated with 40 nM Isoproterenol and illuminated with 430 nm LED for 2 min.

## DISCUSSION

This study was undertaken because we obtained conflicting results from very similar experiments conducted on two different FLIM microscopes. Because our studies require collection of long timelapse series of FLIM images from the same cells, each of these setups had been previously optimized to achieve highest speed and photon efficiency in its class^6,19^. It therefore was surprising that response dynamics appeared so different. Our results show that differences in blue light excitation intensity are causative, not by affecting performance of the FRET sensor or by affecting turnover of intracellular cAMP, but rather by photo-oxidizing all three β-adrenergic agonists (Isoproterenol, Norepinephrine and Adrenaline) used in this study.

Historically, confocal point scanning microscopes have not been regarded as the first choice to circumvent phototoxicity. First, the confocal pinhole significantly diminishes photon efficiency. Moreover, bleaching and phototoxicity are both strongly non-linear^33^, and thus concentration of the excitation light in a single focal spot for a short time (pixel dwell time in the order of a few µs) would likely be unnecessarily harsh to the cells. In widefield microscopes the excitation is spread evenly throughout the field of view and over a prolonged integration time (0.2-2 s / FLIM image), supposedly permitting cells to better cope with radicals using their internal scavenging systems. This simple fact has been the rationale for the development of widefield deconvolution microscopes as alternative for confocal point scanners to study photosensitive processes such as mitosis^21,34^.

Unexpectedly, however, our results show unequivocally that in these experiments the data acquired with the confocal TCSPC system are more reliable. This is most likely because the TCSPC paradigm appears to be considerably more photon efficient than fdFLIM or even siFLIM to arrive at reliable pixel lifetimes, a quality summarized by Gerritsen and colleagues in the figure of merit^35,36^. Thus, typical excitation powers measured at the preparation plane were about 45 µW for TCSPC, contrasting with 900 µW for the fdFLIM system to achieve comparable S/N ratio.

The observed difference held true for different cell lines (Fig. S6) and for three different β1 agonists (Fig. 7). Blue and UV light exposure is well established to induce ROS^11,13,22^ and in literature, there is ample precedence for light sensitivity in cAMP metabolism. Akaike and colleagues discuss that blue light mediated ROS or Reactive Nitrogen Species (RNS) production could lead to formation of nitrated cyclic nucleotides (e.g. 8-nitro-cGMP)^37,38^, which in turn could activate PDE2 and PDE3, both of which are known to be upregulated by cGMP production to increase cAMP breakdown^39,40^. In addition, PDE isoforms directly activated by blue light have been reported in bacteria^41,42^, and blue-light modulation of AC activity has also been described^43–46^. We therefore set out to asses which stage in the biological process is most sensitive for blue light excitation. However, in a variety of experiments, we found no evidence for blue-light induced changes in cAMP metabolism. We also note that in pilot experiments carried out with a cGMP FRET sensor^47^, we never obtained any evidence for blue-light induced cGMP formation in Cos7 and HeLa cells (data not shown). The possibility that blue light/ROS inactivates the β-receptor faster, thus rendering the responses more transient, is also ruled out by the experiments in Fig. 6B and Fig. S4.

Thus, however unlikely at first glance, we had to consider that blue light interferes with activation of β-receptors by all three agonists, IsoP, NE and AR. Indeed, a literature search revealed that Riboflavin (Vitamin B2), which is present at 1 µM in DMEM formulations, can form radicals under the influence of blue light which can specifically attack IsoP so as to form N-Isopropylaminochrome and inactivate binding to the receptor^25,26^. However, we found no literature on a similar sensitivity of NE or AR to Riboflavin radicals, and importantly, our experiments were intentionally carried out in FluoroBrite (FB) medium, which is devoid of Riboflavin and specifically formulated to reduce phototoxicity and autofluorescence. Yet, the experiments described in Figs 7 to 10 show that this hypothesis is correct. In the absence of manufacturers data on the composition of FluoroBrite medium we had to resort to extensive mass spectrometry analysis and time-consuming FRET experiments to reveal that indeed the FB medium is the culprit. Our data show that Folic acid (Vitamin B9), which is also an ingredient of DMEM mediums, can also inactivate all three β-receptor agonists under the influence of blue light.

These data also strongly emphasize the crucial importance of exact knowledge on the molecular composition of reagents, buffers and media. The practice that recipes are kept secret and kits are offered as black-boxes solely for commercial benefit is simply not in accordance with good scientific practice which aims at a full, exact and reproducible description of experiments. More generally, this study should caution researchers who perform live-cell time-lapse experiments to be aware about sample health, phototoxicity and image acquisition routines. Subtle phototoxicity effects, even though invisible morphologically, can still alter kinetic and dynamic responses of cells and thus, biological outcome. The drugs that we have tested in this study (IsoP, NE, AR) are very commonly used catecholamines, routinely used to study the signaling pathways they stimulate using biosensors and live cell imaging^48,49^. Therefore it is crucial that we understand the effect of the imaging regime used in our studies.

## Supporting information

Supplemental data - Radical differences between two FLIM microscopes affect interpretation of cell signaling dynamics

## Acknowledgements

We are indebted to dr. B. Ponsioen (Center of Molecular Medicine at the University Medical Center Utrecht), prof. dr. H. Gerritsen (Utrecht University) and dr. O. Kukk (Solis BioDyne) for insightful discussions and to dr. R. Harkes (NKI BioImaging facility) for preparing the ImageJ plugin to import and interpret fdFLIM data. Experiments in Fig. 6A were performed by master student Enrico Santini (Vrije University, Amsterdam). Dr. O. Kukk is also acknowledged for initial expert assistance with fdFLIM experiments.

This study was supported by funding from NWO (TTW 14691 to K. Jalink). Research at the Netherlands Cancer Institute is supported by institutional grants of the Dutch Cancer Society and of the Dutch Ministry of Health, Welfare and Sport.

## Author contributions

Conception of the study: S.M. and K.J., design and execution of FLIM experiments: S.M. and K.J.; preparation of molecular constructs, construction of stable cell lines and cell culturing: J.K. and S.M., ImageJ scripts for automatic cell segmentation and extraction of single-cell FLIM traces: B.vd.B.; data analysis, statistics and preparation of figures: S.M., B.vd.B., F.E.O. and K.J.; ELISA assays: S.M.; mass spectrometry experiments: F.E.O.; preparation of the manuscript: K.J. and S.M. All authors provided critical input during finalizing of the manuscript.

## MATERIALS AND METHODS

### Cell culture and generation of stable lines

HeLa cells (CCL-2), Cos7 cells (CRL-1651), Hek cells (CRL-1573), MelJuso cells (HTB-68), MCF-7 cells (HTB-22) and A431 cells (CRL-1555) were all cultured in DMEM (Gibco, 31966-021) supplemented with 10% FCS (Gibco, 10270). Stable cell lines were created either using Epac-S^H250^ biosensor or using Epac-S^H201^ biosensor^18^. Epac-S^H250^ is a variant of the Epac-S^H201^ biosensor^18^ with an added P2A-PA-mCherry-HNS and neomycin resistance gene instead of puromycin resistance. For that, transfection of each cell type was performed with the *Tol2* transposon system^18^. For transfection two plasmids are used: a cDNA with the transposase sequence and another cDNA with the following elements: Tol2 sequence, a promoter, the neomycin resistance gene (H250) or puromycin resistance gene (H201), the gene encoding for Epac-S^H250^ or Epac-S^H201^ and a second *Tol2* sequence.

The different cell types were seeded on 6-well plates at approximately 10% density and transfected the next day. 1 μg of both plasmids was mixed with 6 μl FuGENE reagent (Promega E269A) in 200 μl serum free DMEM and incubated for 30 min before adding the transfection mix to the cells. The cells were further incubated for 48 h and subsequently subjected to Gentamycin (G418) selection (1 μg/ml, Roche, G418-RO). After 4 days cells were sorted on a fluorescence-activated cell sorter (FACS) based on mTurq2 fluorescence intensity.

### Compounds and Mediums used in the experiments

**Table 1:** List of compounds and mediums used in FRET-FLIM experiments. HBS was prepared in the lab and contained (in mM): 140 NaCl, 5 KCl, 1 MgCl_2_, 1 CaCl_2_, 10 HEPES, adjusted to pH 7.3 using NaOH. 10 mM glucose was added just before use.

| Name | Supplier | Final concentration |
| --- | --- | --- |
| Isoproterenol | Sigma Aldrich, PHR2722 | 40 nM |
| Forskolin | Sigma Aldrich, F3917 | 25 µM |
| Norepinephrine | Sigma Aldrich, A7257 | 200 nM |
| Adrenaline | Sigma Aldrich, Y0000882 | 250 nM |
| Prostaglandin-E1 | Sigma Aldrich, 538903 | 200 nM |
| Ascorbic Acid | Sigma Aldrich, A92902 | 100 µM |
| Riboflavin | Sigma Aldrich, 555682 | See text |
| DMEM | Gibco, 31966-021 | N/A |
| FluoroBrite | Gibco, A18967-01 | N/A |
| Hepes Buffered Saline (HBS) | made freshly |  |

### Microscopy assay on both setups

To monitor the production and breakdown of cAMP in real time, the donor (mTurquoise2) fluorescence lifetime of the Epac-based FRET biosensor was measured by FLIM. Our FLIM sensor features a tandem dark (i.e., non-emitting) Venus acceptor which allows recording a large part of the donor emission spectrum while minimizing contamination of the signal with acceptor emission^20^.

#### Confocal TCSPC setup (Leica Stellaris and FALCON)

FLIM experiments were carried out using a Leica TCS-SP8 FALCON and Leica Stellaris 8 FALCON confocal FLIM microscope using LAS-X version 3.5 software (SP8) and 4.6.0.27096 software (Stellaris 8). The microscope was equipped with a humidified incubator with 5% CO_2_ at 37 °C. Cells were excited with the 440 nm pulsed diode laser (PicoQuant; SP8) or the 440 nm line of the white-light laser (Stellaris), and photon arrival times were recorded with one HyD detector, covering the mTurquoise emission spectrum (467–500 nm). Cells were grown on 24 mm coverslips (Marienfeld, 0117640), mounted in a steel Leiden incubation chamber and inserted on the microscope stage. A single randomly selected field of view with ∼ 200–600 cells per Field-of-View (FOV) was imaged at 5 s timelapse rate. In all experiments, the position of the focal plane was actively stabilized using the Leica Auto Focus Control (AFC) to prevent focal drift or focus artifacts from pipetting of stimuli.

**Table 2:** Settings on the TCSPC/Confocal setup for imaging.

|  |  |
| --- | --- |
| Acquisition mode | FLIM (xyt) |
| Frame repetitions | 1 |
| FLIM channel assignment | HyD2 (467 nm-500 nm) |
| Format | 512 x 512 |
| Speed | 400 lines/s |
| Zoom factor | 0.75 |
| Size of the FOV | 774 µm x 774 µm |
| Pixel size | 1.51 µm x 1.51 µm |
| Optical section | 6.8 µm |
| Pixel dwell time | 3.16 µs |
| Pinhole | 5.6 airy unit |
| Time interval | 5 second |
| Laser settings | 440 nm, 25% |
| Objective | HC PL APO CS2 20x/Dry (NA = 0.75) |
| Framerate | 0.775/s |

#### Widefield fdfLIM setup (Lambert Instruments)

Frequency-domain FLIM measurements were performed with an optimized MEMFLIM-based camera system^6,9,50^ and recorded using LI-FLIM software version 1.4.0 (Lambert Instruments, Groningen, the Netherlands), attached to a Leica DM-IRE2 widefield microscope. Prior to any experiments on the fdFLIM setup, a reference was recorded with Rhodamine 6G (5 µM dissolved in deionized water; Lifetime 3.93 ns). Calibration of the system was performed by placing the reference on a 40x oil immersion objective (NA = 1.25) and focusing until maximum intensity of the reference was reached using filter cube 3, see below, unless otherwise mentioned. Hereafter, 5 images were taken of the reference (200 ms) at 37 °C and only the last image was saved as actual reference for the experiment so as to minimize the effects of thermal drift in the camera. Further settings were:

**Table 3:** Settings on the fdFLIM/Widefield setup for imaging. Note that for the data in Fig. 1C only, an extra correction was applied: lifetimes were multiplied by 0.77 to compensate for a known error in hardware.

|  |  |
| --- | --- |
| Exposure time | 200 ms/frame |
| Custom phase order (deg) | 0, 120, 240, 180, 300, 60, 270, 30, 150, 90, 210, 330 |
| Camera gain | 2 |
| Modulation frequency (MHz) | 40 |
| Number of phases | 12 |
| LED | 1 Watt – 442 nm LED |
| Toggel register | First and Second |
| Filter cube 2 | ET500/20; 515LP; 535/30 |
| Filter cube 3 | ET438/28; 470LPXR; 465LP |
| Time interval | 5 seconds |

#### Image analysis/cAMP dynamics analysis for fdFLIM and TCSPC

##### Confocal TCSPC setup (Leica Stellaris, FALCON)

The recorded TCSPC photon arrival time histograms showed multi-exponential decay, suggesting a superposition of (at least) two FRET states. Therefore, he photon arrival times were fitted to a double-exponential reconvolution function with fixed lifetimes *τ*_i_ of 0.6 ns and 3.4 ns, representing a high-FRET state and low-FRET state, respectively, using Leica LAS X software. The resulting two timelapse images contained for every pixel the amplitudes (*A*_i_) of these two components. They were exported from LAS X as *ImageJ TIF* files with 0.001 ns per gray value. The *TIF* files were then further processed and analyzed with a custom ImageJ macro in Fiji^51^. In order to relate to conventionally reported (average) lifetime values we map the ratio of the components back to the original 0.6 – 3.4 ns scale, resulting in a weighted mean lifetime value: 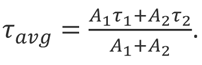 The summed intensity image was used as input for cell segmentation using the deep learning network Cellpose^52^. The pre-trained model ‘cyto2’ provided excellent segmentation for our images. The resulting label maps were used to calculate intensity-weighted single-cell lifetime traces, where each pixel belonging to a cell is weighted with its intensity. The data from these traces were exported as.csv files and run through a R script for visualization and statistical analysis.

##### Widefield fdfLIM setup (Lambert Instruments)

FLIM data from the widefield fdFLIM setup (Lambert Instruments) is saved in the *FLI* format. Reference dye.*FLI* file and sample .*FLI* files were opened and processed with the same script in Fiji, but herein a custom plugin (https://github.com/rharkes/fdFLIM)^18^ was used that opens the .*FLI* files and calculates the lifetimes from both phase and modulation for a single timepoint. The phase lifetime images were concatenated into timelapse series, before they were further processed in the same manner as described above.

### Data visualization and Statistical analysis

A custom R script written in RStudio (version 2023.03.0) was used for data visualization and statistical analysis (https://github.com/Jalink-lab/FLIM-timelapse-traces). Briefly, the script runs through the csv file which contains the lifetime data of every cell imaged per time-point. Following determination of baseline lifetime values for each cell from the first 10 time points, the script allows setting criteria for inclusion of each individual cell based on outlier rejection and variance of time points within the baseline. Typically, between 1 and 15% of cells were rejected, often because of poor segmentation and/or detached cells that affect the validity of the data.

Since typical FOV contain up to 700 individual cells, we choose to plot the traces of 20 individual cells, picked randomly from all cells that pass inclusion criteria. To still fully capture the range and variability of the data, we also included a summary dotplot of the relevant data from all included cells, displayed next to the traces and at the same Y axis scale. Dotplots also show boxplots with median value (horizontal black line), middle 50% of values (boxes) and 1.5 times the interquartile range (whiskers). Full width half maximum (FWHM) in Fig. 1, 3 and 6 were estimated by averaging the results obtained from eye-balled quantitation from 200 cells (confocal) and from 50 cells (widefield) each. The figures were assembled using Inkscape 1.1. All traces displayed in figures 1-3, 5-8 and 10 are established in this manner.

### Power measurements

The power of the illumination light from both setups and also the different LEDs used were measured using a power meter (Newport, model:1830-C) using a 1cm^2^ detector held directly in front of the objective. For detection of LED power, the detector was held at a distance corresponding to the illumination distance between LED and cells. Measured powers are wavelength-corrected.

### Different LEDs used

**Table 4:**
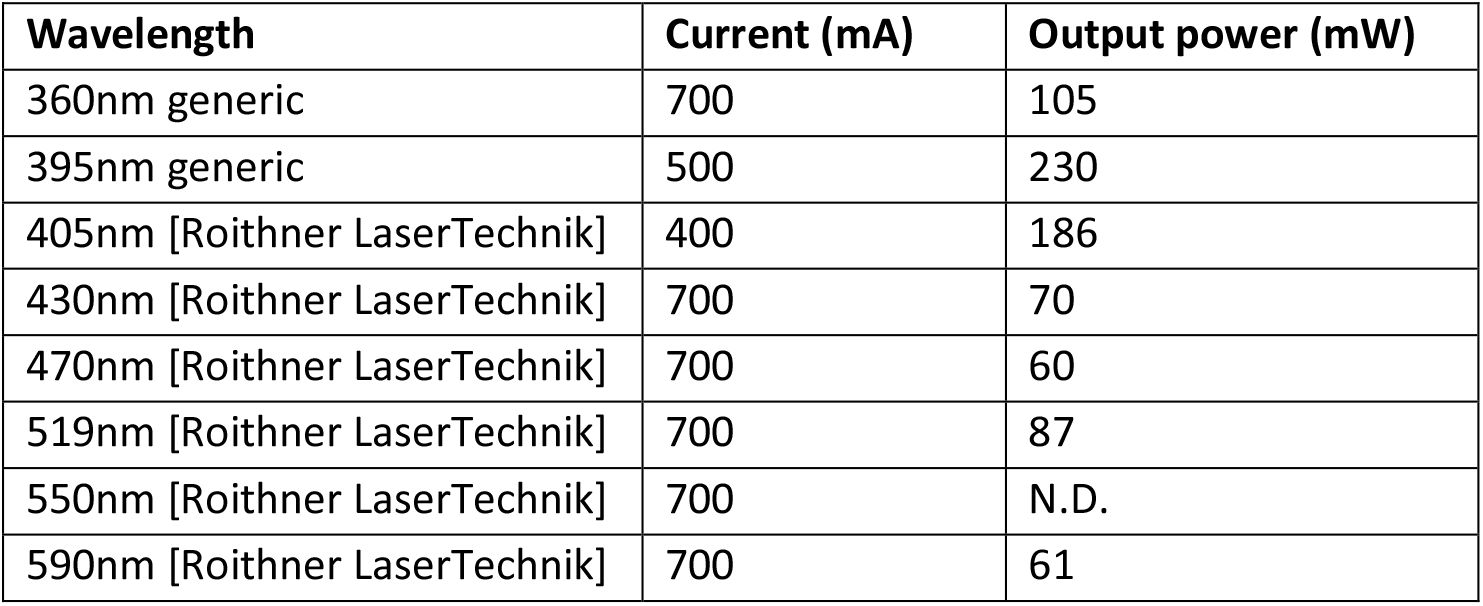
Electrical current and optical power of LEDs used.

#### cAMP ELISA assay

cAMP ELISA assay was performed using a Cyclic AMP XP^®^ Assay Kit (#4339, Cell Signaling Technology) according to the manufacturers protocol. Cells were seeded in a 6 well plates and grown overnight. All experiments were performed in biological duplicates. The following samples were tested :

- Control, untreated cells (Cos7H250 and Cos7 WT)
- Cells treated with 40nM Isoproterenol (Cos7H250 and Cos7WT)
- Cells treated with 40nM Isoproterenol and subjected to 430nm LED light exposure for 2min (Cos7H250 and Cos7 WT)

Note that where cAMP levels outrange the maximum value for the ELISA assay, the max values of the calibration curve (120 nM) are shown instead.

#### ROS dye assay

The ROS dye assay was performed using Di(Acetoxymethyl Ester) (6-Carboxy-2’,7’-Dichlorodihydrofluorescein Diacetate) ROS dye (Invitrogen, D23844), a fluorogenic dye that is converted to 6-CarboxyFluorescein, which is fluorescent (λ_ex_494 nm, λ_em_518 nm) upon exposure to ROS.

For the assay, Cos7 WT cells were seeded on coverslips in a 6 well plate and grown overnight. On the day of the experiment, the cells were washed twice with PBS and 2 ml of FluoroBrite (FB) medium with 0.8 µL of the stock dye (final concentration of 10 µM) was added to each of the 6 wells. The 6 well plate was covered with aluminum foil to avoid exposure to light and incubated for 45 min. Following that, the incubation mix was pipetted off and the cells were washed with 2 ml of FB medium. Coverslips where mounted in Leiden holders, put on the microscope stage and 3 different FOVs were imaged using the widefield fdFLIM setup at using standard settings (see widefield fdFLIM setup) except that Filter cube 2 was used. Then, the same coverslip is used to image 3 different FOVs using the TCSPC confocal setup at settings described above (laser 488 nm is used at 2% excitation power).

#### Caged cAMP assay

For stimulation by photo-release of caged cAMP, cells were treated with DMNB-caged cAMP (4,5-Dimethoxy-2-Nitrobenzyl Adenosine 3’,5’-Cyclicmonophosphate, Molecular Probes, D1037) for at least 30 min prior to imaging at a final concentration of 1 mM. Uncaging was with a 200 ms UV pulse from the Leica EL6000 lamp (4 mW, approximately 400 mW/cm^2^ in our experimental setup with the 20×, 0.75 NA dry objective) using a DAPI_LP filter cube (380/40 nm, 405 nm dichroic) which was inserted in the confocal excitation path to enable UV illumination and confocal FLIM recording simultaneously.

#### Mass-spec assays

LC-MS measurements were performed on a system equipped with a Waters 2795 Seperation Module (Alliance HT), Waters XSelect^TM^ CSH^TM^ C18 (4.6 x 100 mm, 5 μm) and Waters LCT Premier XE Mass Spectrometer. Samples were run (column temperature= 40°C, flow= 0.8 mL/min) using 2 mobile phases: A= 1% CH_3_CN and 0.1% formic acid in water and B= 1% water and 0.1% formic acid in CH_3_CN. After injection of the sample (50 μL), an isocratic run of 5%B for 6 min was performed first followed by the following gradient: 0.0 – 1.4 min: 5 – 30%B, 1.4 – 4.2 min: 30 – 80%B, 4.2 – 4.4 min: 80 – 95%B, 4.4 – 5.0 min: 95%B, 5 – 5.3 min: 95 – 5%B, 5.3 – 6 min: 5%B. Data processing was performed using Waters MassLynx Mass Spectrometry Software 4.1.

## Data availability

All data can be found on Zenodo repository: https://doi.org/10.5281/zenodo.8263078

Custom software can be found on GitHub: https://github.com/Jalink-lab/FLIM-timelapse-traces

## References

1. Trebak, M. & Kinet, J.-P. Calcium signalling in T cells. Nat. Rev. Immunol. 19, 154–169 (2019).

2. Berridge, M. J. Module 6: Spatial and Temporal Aspects of Signalling. Cell Signal. Biol. 6, csb0001006 (2014).

3. Peng, G. E., Pessino, V., Huang, B. & von Zastrow, M. Spatial decoding of endosomal cAMP signals by a metastable cytoplasmic PKA network. Nat. Chem. Biol. 17, 558–566 (2021).

4. Zaccolo, M., Zerio, A. & Lobo, M. J. Subcellular Organization of the cAMP Signaling Pathway. Pharmacol. Rev. 73, 278–309 (2021).

5. van Rheenen, J., Langeslag, M. & Jalink, K. Correcting Confocal Acquisition to Optimize Imaging of Fluorescence Resonance Energy Transfer by Sensitized Emission. Biophys. J. 86, 2517–2529 (2004).

6. Raspe, M. et al. siFLIM: single-image frequency-domain FLIM provides fast and photon-efficient lifetime data. Nat. Methods 13, 501–504 (2016).

7. Wu, H.-M. et al. Widefield frequency domain fluorescence lifetime imaging microscopy (FD-FLIM) for accurate measurement of oxygen gradients within microfluidic devices. Analyst 144, 3494– 3504 (2019).

8. Elder, A. D. et al. The application of frequency-domain Fluorescence Lifetime Imaging Microscopy as a quantitative analytical tool for microfluidic devices. Opt. Express 14, 5456–5467 (2006).

9. Specifications. Lambert Instruments https://www.lambertinstruments.com/toggel-specifications.

10. Shen, Y. et al. A genetically encoded Ca2+ indicator based on circularly permutated sea anemone red fluorescent protein eqFP578. BMC Biol. 16, 9 (2018).

11. Laissue, P. P., Alghamdi, R. A., Tomancak, P., Reynaud, E. G. & Shroff, H. Assessing phototoxicity in live fluorescence imaging. Nat. Methods 14, 657–661 (2017).

12. Kiepas, A., Voorand, E., Mubaid, F., Siegel, P. M. & Brown, C. M. Optimizing live-cell fluorescence imaging conditions to minimize phototoxicity. J. Cell Sci. 133, jcs242834 (2020).

13. Icha, J., Weber, M., Waters, J. C. & Norden, C. Phototoxicity in live fluorescence microscopy, and how to avoid it. BioEssays 39, 1700003 (2017).

14. Jonkman, J. & Brown, C. M. Any Way You Slice It—A Comparison of Confocal Microscopy Techniques. J. Biomol. Tech. JBT 26, 54–65 (2015).

15. Lucius, R., Mentlein, R. & Sievers, J. Riboflavin-Mediated Axonal Degeneration of Postnatal Retinal Ganglion Cells In Vitro is Related to the Formation of Free Radicals. Free Radic. Biol. Med. 24, 798–808 (1998).

16. Bogdanov, A. M., Kudryavtseva, E. I. & Lukyanov, K. A. Anti-Fading Media for Live Cell GFP Imaging. PLOS ONE 7, e53004 (2012).

17. Edwards, A. M., Silva, E., Jofré, B., Becker, M. I. & De Ioannes, A. E. Visible light effects on tumoral cells in a culture medium enriched with tryptophan and riboflavin. J. Photochem. Photobiol. B 24, 179–186 (1994).

18. Harkes, R. et al. Dynamic FRET-FLIM based screening of signal transduction pathways. Sci. Rep. 11, 20711 (2021).

19. Application Note: SP8 FALCON: a novel concept in fluorescence lifetime imaging enabling video-rate confocal FLIM.

20. Klarenbeek, J., Goedhart, J., Batenburg, A. van, Groenewald, D. & Jalink, K. Fourth-Generation Epac-Based FRET Sensors for cAMP Feature Exceptional Brightness, Photostability and Dynamic Range: Characterization of Dedicated Sensors for FLIM, for Ratiometry and with High Affinity. PLOS ONE 10, e0122513 (2015).

21. Dixit, R. & Cyr, R. Cell damage and reactive oxygen species production induced by fluorescence microscopy: effect on mitosis and guidelines for non-invasive fluorescence microscopy. Plant J. 36, 280–290 (2003).

22. Douthwright, S. & Sluder, G. Live Cell Imaging: Assessing the Phototoxicity of 488 and 546 nm Light and Methods to Alleviate it. J. Cell. Physiol. 232, 2461–2468 (2017).

23. Figueroa, D., Asaduzzaman, M. & Young, F. Real time monitoring and quantification of reactive oxygen species in breast cancer cell line MCF-7 by 2′,7′–dichlorofluorescin diacetate (DCFDA) assay. J. Pharmacol. Toxicol. Methods 94, 26–33 (2018).

24. Eruslanov, E. & Kusmartsev, S. Identification of ROS Using Oxidized DCFDA and Flow-Cytometry. in Advanced Protocols in Oxidative Stress II (ed. Armstrong, D.) 57–72 (Humana Press, 2010). doi:10.1007/978-1-60761-411-1_4.

25. Massad, W. A., Bertolotti, S. & Garcia, N. A. Kinetics and Mechanism of the Vitamin B2– sensitized Photooxidation of Isoproterenol¶. Photochem. Photobiol. 79, 428–433 (2004).

26. Massad, W. A., Marioli, J. M. & García, N. A. Photoproducts and proposed degradation pathway in the riboflavin-sensitised photooxidation of isoproterenol. Pharm. 61, 1019–1021 (2006).

27. Insińska-Rak, M. & Sikorski, M. Riboflavin Interactions with Oxygen—A Survey from the Photochemical Perspective. Chem. – Eur. J. 20, 15280–15291 (2014).

28. FluoroBrite^TM^ DMEM. https://www.thermofisher.com/order/catalog/product/A1896701.

29. Margineanu, M. B. et al. Semi-automated quantification of living cells with internalized nanostructures. J. Nanobiotechnology 14, 4 (2016).

30. 11995 - DMEM, high glucose, pyruvate - NL. https://www.thermofisher.com/nl/en/home/technical-resources/media-formulation.9.html.

31. Lowry, O. H., Bessey, O. A. & Crawford, E. J. PHOTOLYTIC AND ENZYMATIC TRANSFORMATIONS OF PTEROYLGLUTAMIC ACID. J. Biol. Chem. 180, 389–398 (1949).

32. Baibarac, M., Smaranda, I., Nila, A. & Serbschi, C. Optical properties of folic acid in phosphate buffer solutions: the influence of pH and UV irradiation on the UV-VIS absorption spectra and photoluminescence. Sci. Rep. 9, 14278 (2019).

33. Hoebe, R. A. et al. Controlled light-exposure microscopy reduces photobleaching and phototoxicity in fluorescence live-cell imaging. Nat. Biotechnol. 25, 249–253 (2007).

34. Rieder, C. L. & Khodjakov, A. Mitosis Through the Microscope: Advances in Seeing Inside Live Dividing Cells. Science 300, 91–96 (2003).

35. Gerritsen, H. C., Asselbergs, M. a. H., Agronskaia, A. V. & Van Sark, W. G. J. H. M. Fluorescence lifetime imaging in scanning microscopes: acquisition speed, photon economy and lifetime resolution. J. Microsc. 206, 218–224 (2002).

36. de Grauw, C. J. & Gerritsen, H. C. Multiple Time-Gate Module for Fluorescence Lifetime Imaging. Appl. Spectrosc. 55, 670–678 (2001).

37. Akaike, T., Nishida, M. & Fujii, S. Regulation of redox signalling by an electrophilic cyclic nucleotide. J. Biochem. (Tokyo) 153, 131–138 (2013).

38. Sawa, T. et al. Formation, signaling functions, and metabolisms of nitrated cyclic nucleotide. Nitric Oxide Biol. Chem. 34, 10–18 (2013).

39. Pavlaki, N. & Nikolaev, V. O. Imaging of PDE2-and PDE3-Mediated cGMP-to-cAMP Cross-Talk in Cardiomyocytes. J. Cardiovasc. Dev. Dis. 5, 4 (2018).

40. Polito, M. et al. The NO/cGMP pathway inhibits transient cAMP signals through the activation of PDE2 in striatal neurons. Front. Cell. Neurosci. 7, 211 (2013).

41. Tian, Y., Yang, S., Nagel, G. & Gao, S. Characterization and Modification of Light-Sensitive Phosphodiesterases from Choanoflagellates. Biomolecules 12, 88 (2022).

42. Miki, N., Baraban, J., Keirns, J., Boyce, J. & Bitensky, M. Purification and properties of the light-activated cyclic nucleotide phosphodiesterase of rod outer segments. J. Biol. Chem. 250, 6320–6327 (1975).

43. Iseki, M. et al. A blue-light-activated adenylyl cyclase mediates photoavoidance in Euglena gracilis. Nature 415, 1047–1051 (2002).

44. Stierl, M. et al. Light Modulation of Cellular cAMP by a Small Bacterial Photoactivated Adenylyl Cyclase, bPAC, of the Soil Bacterium Beggiatoa♦. J. Biol. Chem. 286, 1181–1188 (2011).

45. Tolentino Collado, J., et al. Unraveling the Photoactivation Mechanism of a Light-Activated Adenylyl Cyclase Using Ultrafast Spectroscopy Coupled with Unnatural Amino Acid Mutagenesis. ACS Chem. Biol. 17, 2643–2654 (2022).

46. Ohki, M. et al. Structural insight into photoactivation of an adenylate cyclase from a photosynthetic cyanobacterium. Proc. Natl. Acad. Sci. 113, 6659–6664 (2016).

47. Calamera, G. et al. FRET-based cyclic GMP biosensors measure low cGMP concentrations in cardiomyocytes and neurons. Commun. Biol. 2, 1–12 (2019).

48. Liu, Z. & Liu, S. A novel fluorescent biosensor for adrenaline detection and tyrosinase inhibitor screening. Anal. Bioanal. Chem. 410, 4145–4152 (2018).

49. Feng, J. et al. A genetically encoded fluorescent sensor for rapid and specific in vivo detection of norepinephrine. Neuron 102, 745–761.e8 (2019).

50. Zhao, Q. et al. MEM-FLIM: all-solid-state camera for fluorescence lifetime imaging. in Imaging, Manipulation, and Analysis of Biomolecules, Cells, and Tissues X vol. 8225 93–103 (SPIE, 2012).

51. Schindelin, J., et al. Fiji: an open-source platform for biological-image analysis. Nat. Methods 9, 676–682 (2012).

52. Stringer, C., Wang, T., Michaelos, M. & Pachitariu, M. Cellpose: a generalist algorithm for cellular segmentation. Nat. Methods 18, 100–106 (2021).

