## Supplemental data - Radical differences between two FLIM microscopes affect interpretation of cell signaling dynamics for "‘Radical’ differences between two FLIM microscopes affect interpretation of cell signaling dynamics"

### Supplementary Data:

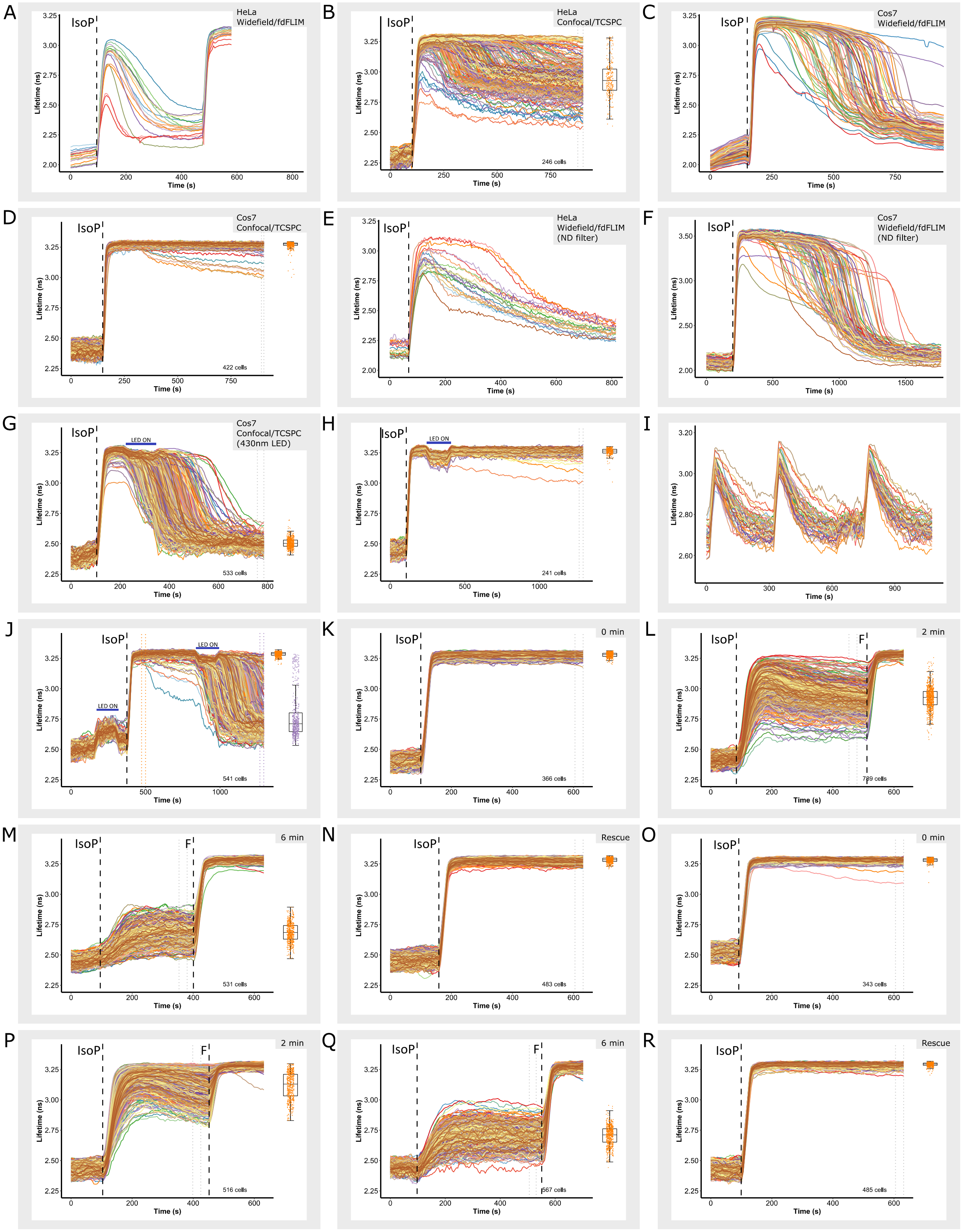


***Figure S1:*** *Cells stably expressing the cAMP FLIM sensor Epac-S$^{H201}$ or Epac-S$^{H250}$ were imaged every 5 s. Following recording of a baseline, cells were challenged with Isoproterenol (IsoP; 40 nM) as indicated, and Forskolin (F; 25 µM) was added to saturate the response for comparison. Shown are FLIM timelapse responses in individual HeLa cells as detected on the* ***(A)*** *widefield fdFLIM setup at standard excitation at 0.9 mW (0.49 W/cm$^2$) using a 435 nm sine-wave modulated LED and* ***(B)*** *on the*

*FALCON TCSPC setup and in individual Cos7 cells detected on the* ***(C)*** *widefield fdFLIM setup and* ***(D)*** *on the FALCON TCSPC system.* ***(E)*** *Responses of HeLa cells to 40nM IsoP recorded using fdFLIM using a ND1 filter in the excitation path to reduce excitation to 0.091 mW (0.05 W/cm$^2$) at the plane of the preparation.* ***(F)*** *Same experiments as in E for Cos7 cells.* ***(G)*** *Transient responses detected by TCSPC when cells were additionally exposed to continuous light from a 430 nm LED for 2 min as indicated.* ***(H)*** *Ascorbic acid reverts the cAMP response to sustained despite exposure to 430 nm LED light for 2 min.* ***(I)*** *Responses to three consecutive doses of cAMP generated by flash photolysis of caged cAMP. After the second dose, cells were exposed to LED light. cAMP decay times were identical for all 3 cases, indicating that PDE activity is unaltered by the light. Note that in this case 470 nm LED was used to prevent slight photolysis of caged cAMP by the 430 nm LED.* ***(J)*** *Blue light fails to evoke response transientness when applied before addition of agonist. Shown are 20 randomly selected traces (see M&M) as well as the response magnitude of all 541 cells taken before (orange boxplot) and after (purple boxplot) LED exposure. Responses to 40 nM Isoproterenol when added from small batches dissolved in 200 µl of FB medium that have been subjected to blue light illumination for* ***(K)*** *0 min,* ***(L)*** *2 min,* ***(M)*** *6 min and* ***(N)*** *for 6 min in the presence of 100 µM Ascorbic acid. Responses to 40 nM Isoproterenol dissolved in 200 µl of HBS containing 18 µM Folic acid in eppendorf vials were subjected to blue light illumination for* ***(O)*** *0min,* ***(P)*** *2min,* ***(Q)*** *6 min* ***(R)*** *for 6 min in the presence of 100 µM Ascorbic acid.*

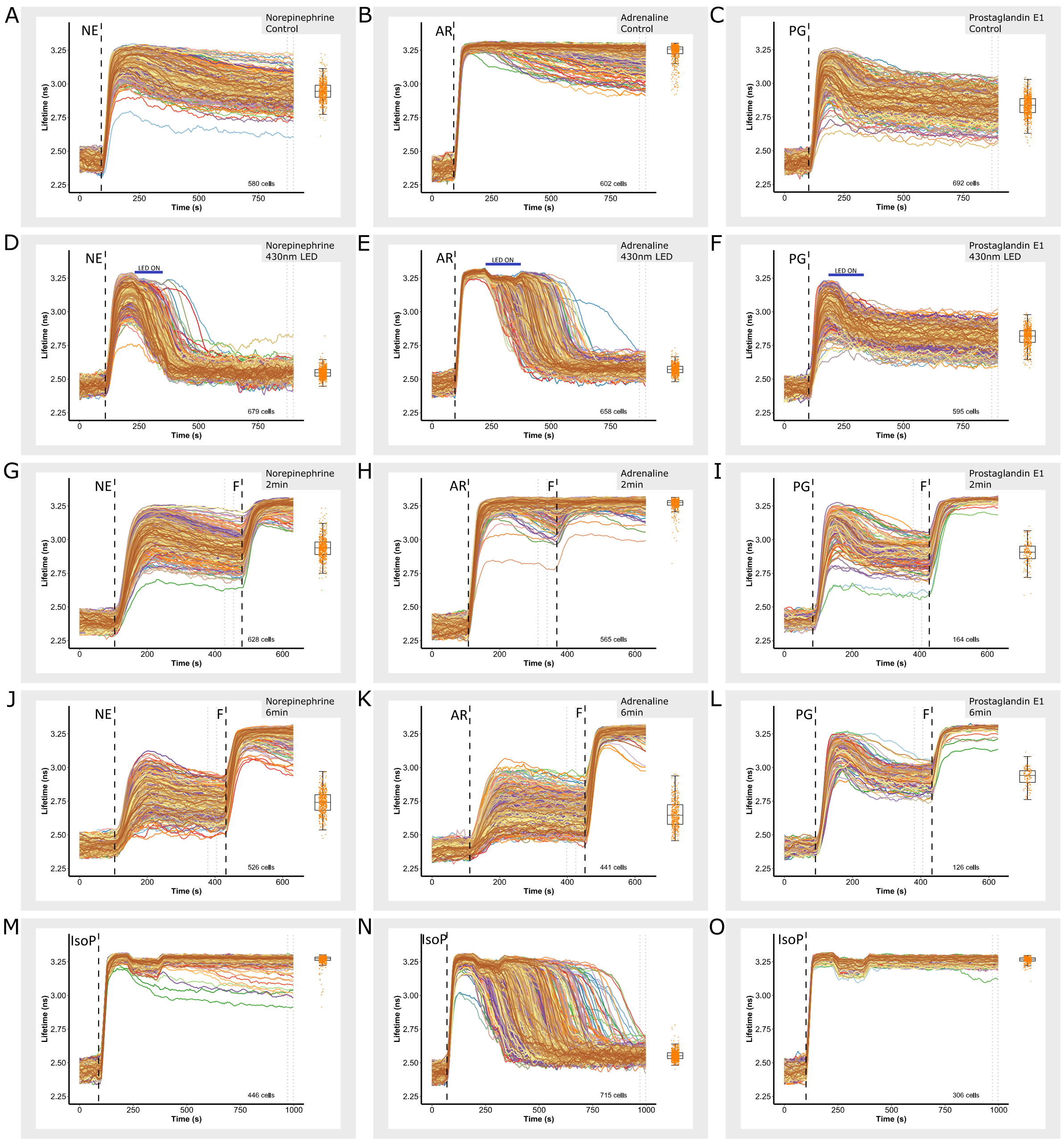


***Figure S2:*** ***(A)*** *Sustained responses of Cos7 cells to 200 nM Norepinephrine (NE)* ***(B)*** *Idem, rendered transient by illumination with 430 nm LED light for 2 min.* ***(C)*** *Sustained responses to 250 nM Adrenaline (AR).* ***(D)*** *Idem, altered to transient upon exposure to 430 nm LED light for 2 min.* ***(E)*** *Typical, partially sustained responses of Cos7 cells to 200 nM Prostaglandin-E1 (PG).* ***(F)*** *Unaltered responses when Cos7 cells are stimulated with 200 nM Prostaglandin-E1 (PG) and illuminated with 430 nm LED light for 2 min.* ***(G,H)*** *Responses to Norepinephrine (200 nM) when added from small batches dissolved in 200 µl of FB medium that have been subjected to blue light illumination for* ***(G)*** *after 2 min pre-exposure to blue light and* ***(H)*** *after 6 min pre-exposure to blue light.* ***(I,J)*** *Idem, responses to Adrenaline (250 nM); shown are* ***(I)*** *after 2 min pre-exposure to blue light and* ***(J)*** *after 6 min pre-exposure to blue light.* ***(K,L)*** *Idem, responses to Prostaglandin-E1 (200 nM)* ***(K)*** *after 2 min pre-exposure to blue light, and* ***(L)*** *after 6 min pre-exposure to blue light.* ***(M)*** *Sustained responses of Cos7 cells in HBS medium only when stimulated with 40 nM Isoproterenol (IsoP) and illuminated with 430 nm LED for 2 min.* ***(N)*** *Transient responses of Cos7 cells in HBS medium containing 45 µM Folic acid when stimulated with 40 nM*

*Isoproterenol (IsoP) and illuminated with 430 nm LED for 2 min.* ***(O)*** *Rescue of transient responses of Cos7 cells in HBS medium containing 45 µM Folic acid and 100 µM Ascorbic acid when stimulated with 40 nM Isoproterenol (IsoP) and illuminated with 430 nm LED for 2 min.*

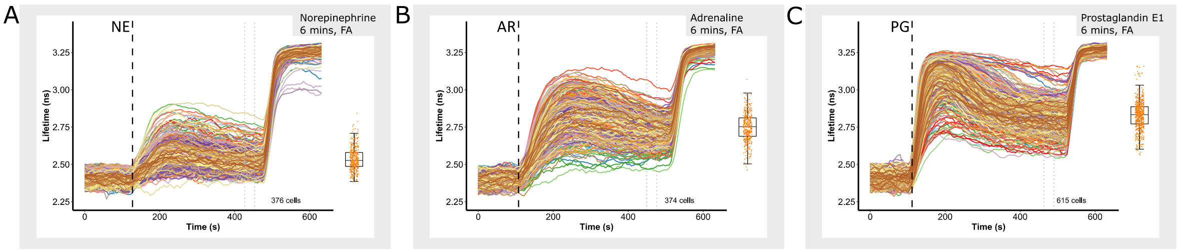


***Figure S3:*** ***(A)*** *200 nM Norepinephrine dissolved in 200 µl of HBS containing 18 µM Folic acid in eppendorf vials were subjected to blue light (430 nm LED) illumination 6 min before use in cell assays.* ***(B)*** *250 nM Adrenaline dissolved in 200 µl of HBS containing 18 µM Folic acid in eppendorf vials were subjected to blue light (430 nm LED) illumination 6 min before use in cell assays.* ***(C)*** *200 nM Prostaglandin-E1 dissolved in 200 µl of HBS containing 18 µM Folic acid in eppendorf vials were subjected to blue light (430 nm LED) illumination 6 min before use in cell assays.*

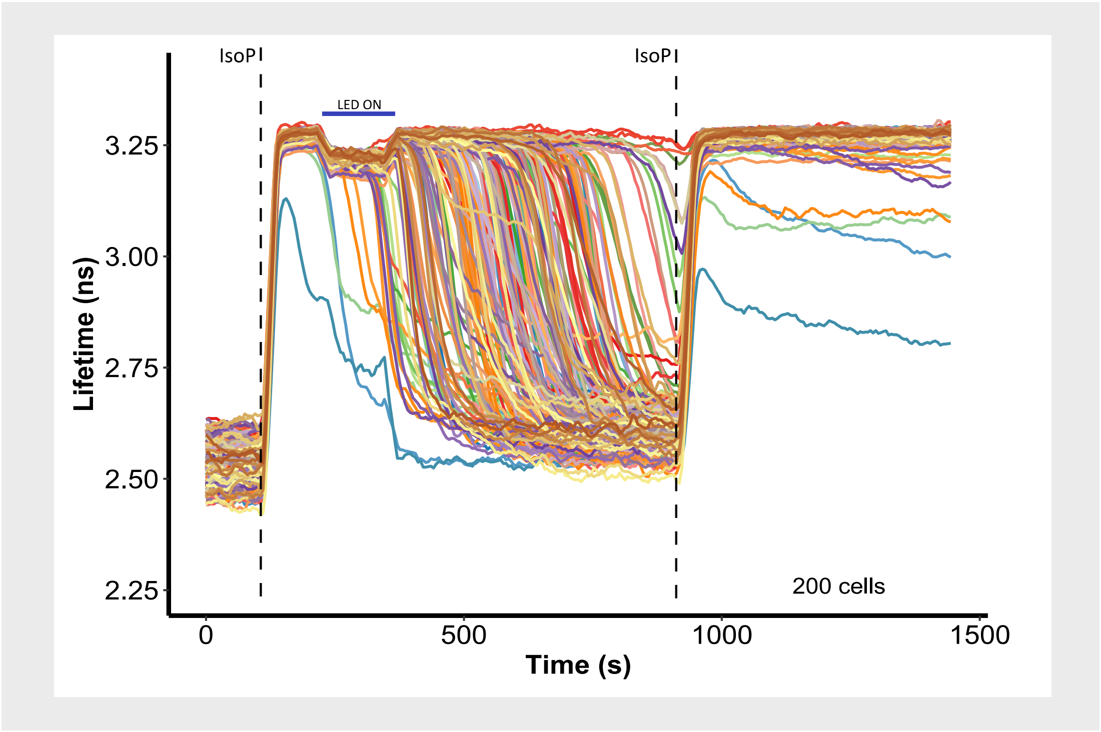


***Figure S4:*** *Cos7 cells stimulated with 40 nM IsoP and subsequently exposed to blue light (430 nm LED) for 2 min, leading to transient cAMP responses. However, a second stimulation with 40 nM IsoP also gives an equally strong increase in cAMP levels, indicated that β-receptors are not saturated and also not inactivated.*

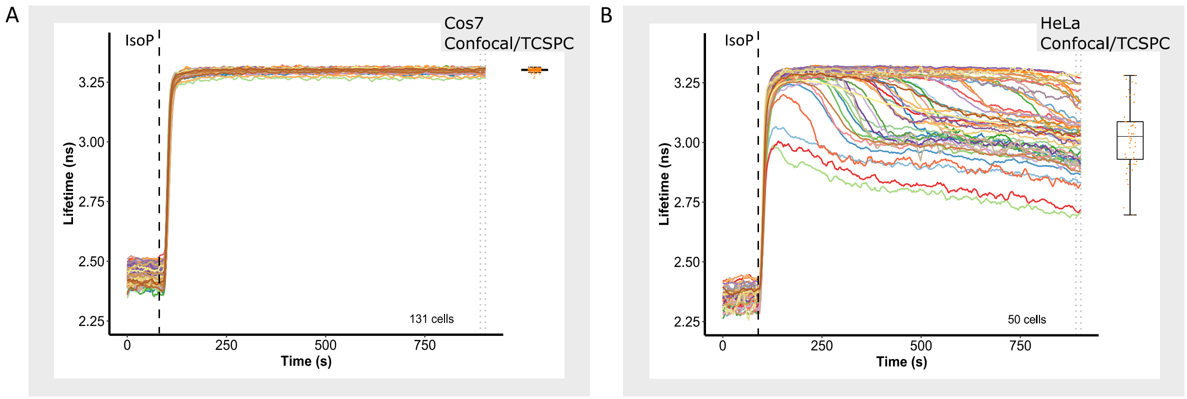


***Figure S5:*** *Cells stably expressing the cAMP FLIM sensor Epac-S$^{H201}$ or Epac-S$^{H250}$ were imaged every 5 s. Following recording of a baseline, cells were challenged with Isoproterenol (IsoP; 40 nM) as indicated. Shown are FLIM timelapse responses in* ***(A)*** *individual Cos7 cells as detected on the FALCON TCSPC setup using a 40x glycerol immersion objective and in* ***(B)*** *individual HeLa cells detected on the on the FALCON TCSPC setup using a 40x glycerol immersion objective.*

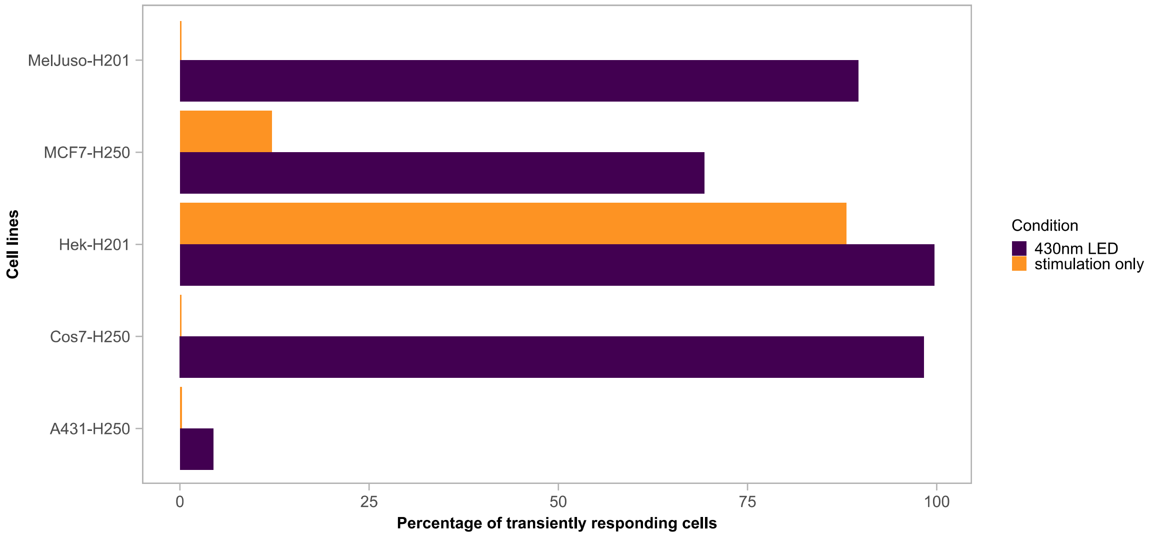


***Figure S6:*** *Different cell lines stably expressing the cAMP FLIM sensor Epac-S$^{H201}$ or Epac-S$^{H250}$ tested for showing transient cAMP responses when exposed to blue light (430 nm LED) for 2 min.*
